# Death of fruit flies at sunset by *Entomophthora muscae* is driven by a host-independent mechanism

**DOI:** 10.1101/2025.06.18.660419

**Authors:** Leslie Torres Ulloa, Dylan Roy, Angie Minichiello, Fosca Becthold, Benjamin de Bivort, Carolyn Elya

## Abstract

Timing in the natural world is a matter of life or death; consequently, nearly all life on Earth has evolved internal circadian clocks. Many behavior-manipulating parasites exhibit striking daily timing, but whether this is clock-driven has remained unclear. Here, we leveraged the laboratory-tractable zombie fruit fly model, *Drosophila melanogaster* infected by the behavior manipulating fungus *Entomophthora muscae*, to tackle this long standing mystery. Using an automated behavioral paradigm, we found that the timing of death of wild-type flies continues to occur with ∼21 hour periodicity in the absence of environmental cues, indicating that death is controlled by a system-intrinsic clock. Experiments with circadian and photoreception mutants revealed that death is independent of host genotype, suggesting that the host does not drive this rhythm and raising the possibility that timing is dictated by *E. muscae*. Transcriptomic analysis of *in vitro* grown fungus revealed that *E. muscae* maintains rhythmic gene expression independent of the fly host that peaks at sunset and has a free-running period of ∼22 hours. Among cycling genes, we identified a transcript encoding a protein with high homology to *white collar-1*, the blue light sensor and core component of the molecular oscillator in the model ascomycete fungus *Neurospora crassa*. Altogether, our findings suggest that *E. muscae* has an endogenous circadian clock. Given that the host genetic background does not appear to dictate the timing of death, our results suggest that the *E. muscae* clock may be the primary driver of this rhythm. Our work supports a model in which a fungal clock influences fly outcomes, suggesting that endogenous parasitic clocks may play a broader role in the temporal coordination of host manipulation.

## Introduction

In nature, much like in comedy, timing is everything. Life on Earth has evolved under the consistent cadence of our planet’s 24-hour rotation, experiencing rhythmic cycles in environmental cues such as light and temperature. In response, organisms across the tree of life have evolved internal timekeeping mechanisms, or circadian clocks, that help align their internal biological processes with daily changes in the environment. Indeed, from microbes to humans, circadian clocks are near universal among living things^1^.

In eukaryotes, circadian clocks generally consist of transcription-translation feedback loops that oscillate with an approximate 24 hour period^2,3^. Circadian clocks are set by environmental cues, termed Zeitgebers (German for “time givers”), that influence circadian gene expression pathways. The most common and often strongest Zeitgeber is light, though others - including temperature^4^, humidity^5^, nutrient availability^6,7^, and even social cues^8^ - can also set the clock.

Though most famous for regulating sleep-wake cycles, circadian clocks influence virtually all aspects of an organism’s biology, including metabolism^9,10^, body temperature^11^, neural plasticity and excitability^12,13^, hormone secretion^14,15^, development^16^, and immunity^17^. Owing to the far-reaching impacts of circadian timing on animal physiology, many parasites have evolved to exploit the predictable timing of host rhythms to enhance their fitness^18,19^. For example, the nematode *Trichuris muris* relies on circadian rhythms in dendritic immune cells of its mouse host to time expulsion into the environment^20^. Similarly, *Plasmodium vivax*, the protozoan that causes recurring malaria, times cell replication with host rhythms which may help avoid peak immune responses or synchronize growth to availability of host resources^21^. *Trypanosoma brucei,* the protist that causes African sleeping sickness, goes one step farther, shortening both behavioral and molecular circadian periods in mice, effectively reprogramming the host clock^22^.

Beyond exploiting and manipulating host rhythms, some parasites have evolved to control host behavior with precise timing^23^. Many examples can be found in the invertebrate world: ants infected with the fungus *Ophiocordyceps* bite onto vegetation and die around solar noon^24^, grasshoppers infected by *Entomophaga grylli* die up to four hours before sunset^25^, nematomorph-infected mantids leap into water at midday^26^, and *Microphallus*-infected snails show aberrant climbing behavior in the early morning^27^. These patterns suggest that time-of-day plays a critical role in maximizing parasite transmission, whether through abiotic conditions or synchronization with other hosts.

While striking, it remains unclear whether host-parasite phenomena that occur with consistent daily timing are the result of proximal environmental cues or internal timekeepers. In some systems, changes in expression of host clock genes during infection and manipulation have led researchers to hypothesize that parasites may manipulate the host’s circadian clock to drive these behaviors^28–30^. However, experimentally demonstrating that the timing is controlled by the host has proven challenging, largely due to the limited genetic tools available.

*Entomophthora muscae* (phylum Zoopagomycota, order Entomophthorales) is a fungal pathogen that infects, manipulates the behavior of, then kills flies. Like many other so-called “zombie-making” parasites, *E. muscae* evokes behavioral changes, and subsequently death, in its host with specific daily timing^31,32^. However, unlike most other such systems, *E. muscae* is amenable to laboratory culture and naturally infects one of the robust genetic models known to biology, the fruit fly *Drosophila melanogaster*^33^. Here, we leverage this laboratory model to address a long-standing question: Is the timing of death by *E. muscae* driven by an internal timekeeper? And if so, does this timekeeper belong to the fly?

## Results

### Death by *E. muscae* follows a time-restricted pattern in the absence of light cues

We first sought to determine whether the specific daily timing of death by *E. muscae-*infected flies persists in the absence of Zeitgebers. To this end, we exposed wild-type Canton-S (hereafter referred to as WT) flies to *E. muscae*, and housed them for 0, 24, 48, 72 or 96 hours with 12 hours light and 12 hours dark (12:12 L:D) light cues before switching to free-running conditions (constant darkness, D:D). As a positive control, we performed the same experiment under 12:12 L:D light cues throughout the experiment. Using an automated behavioral tracking platform^31^, we determined the time when these flies were last observed to move as a proxy for their time of death. We measure this time in Zeitgeber time, with Zeitgeber referring to the light cues that were used to entrain the flies. In light-entrainment paradigms, ZT0 corresponds to the time of day when light turns on; in a 12:12 L:D regime, ZT12 is when lights turn off. Flies were monitored for seven days following exposure to observe all potential deaths by fungus. As seen previously, most deaths occurred on days four and five^31,33^.

We observed that flies die with daily periodicity even in the absence of light cues (**Figure 1A**). Timing of death was most tightly clustered when flies were released into D:D after 96 hours (**Figure 1B**), whose average periodicity was 23.7 hours. When flies were moved to D:D at 72 hours or earlier, the time elapsed between deaths on subsequent days was 20.6 hours (**Figure 1C**). Flies that were housed in D:D from exposure onward (0 L:D) showed the most within day variation among times of death. These data show that time-restricted death by *E. muscae* persists in the absence of light cues, demonstrating that this phenomenon is controlled by an internal time-keeping mechanism.

**Figure 1.**
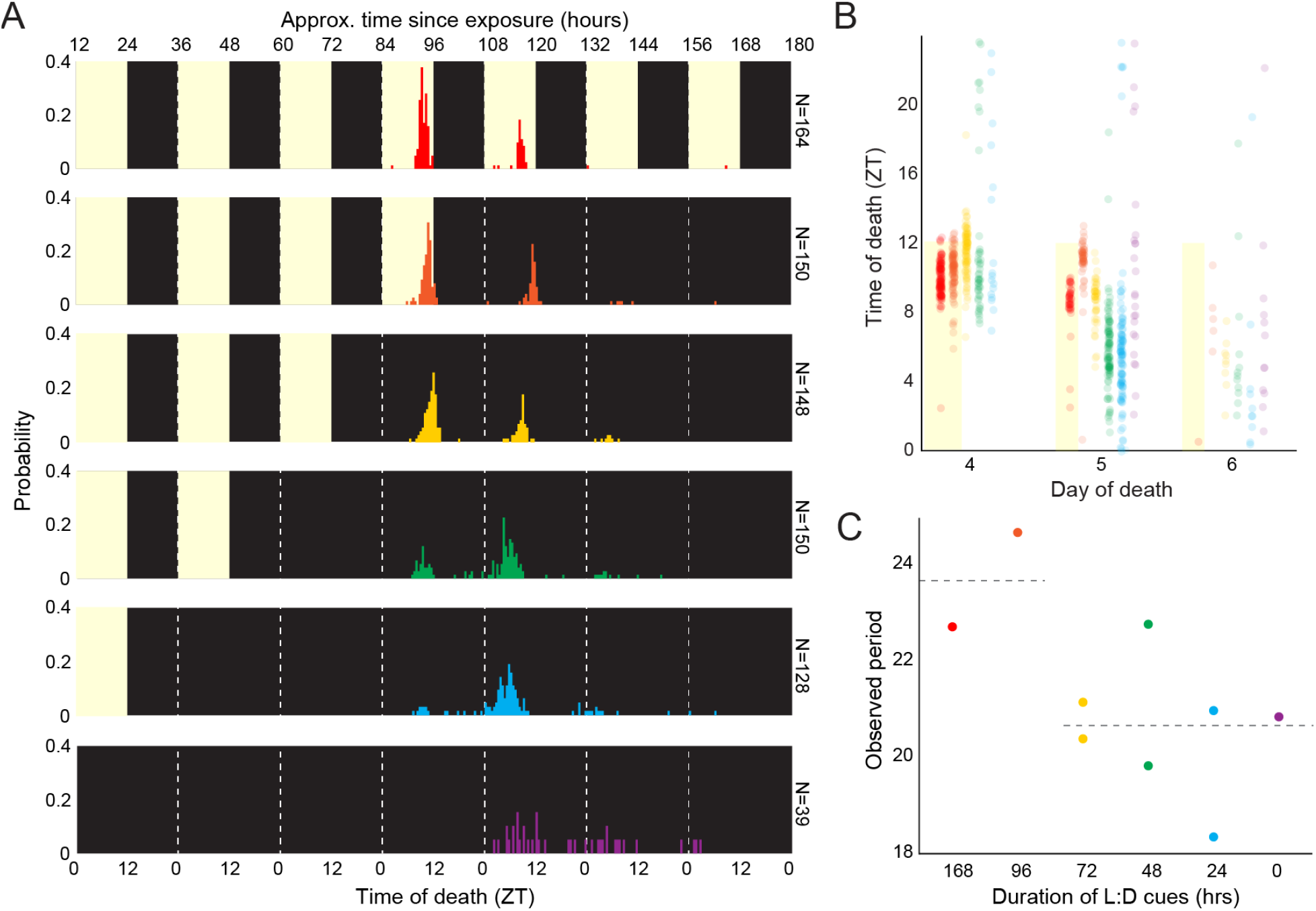
*E. muscae*-infected WT flies die in a gated fashion in the absence of proximal light cues. A) Time of last movement distributions for WT flies exposed to *E. muscae* and housed until varying durations of L:D cues before release into free-running conditions (D:D). Yellow background indicates presence of white light cues during experiment; black indicates darkness. All flies were reared on the same 12:12 L:D cycle (ZT0 = 12:00 AM EST). B) Data from A presented as a scatter plot. C) Observed death period lengths (in hours) based on all observations in A. For every day in which at least 5 flies were observed to die, a mean time of death was calculated for that day; the difference between mean times of death across adjacent days is plotted. Dashed line at left reflects mean observed period for experiments where L:D cues were provided for experiments in which flies had at least one death day with L:D cues (i.e., 168 and 96 hours after *E. muscae* exposure; 23.7 hours); at right, experiments with D:D cues on all death days (i.e., 72, 48, 24, and 0 hours after *E. muscae* exposure), dashed line is mean for these experiments (20.6 hours). Two-tailed t-test shows significant difference between L:D and D:D periods (p=0.0251).

### Success and duration of *E. muscae* infection depends on light

We noticed that the average time of infection lengthened as flies were housed for shorter durations under L:D: flies housed with 24 or 48 hours of L:D cues took significantly longer to die after *E. muscae* exposure (119.4 and 114.7 hours, respectively) compared to 72, 96 and All L:D treatments (106.2, 106.1, and 103.2 hours, respectively) (**Figure 1 - S1A**). Flies kept in darkness throughout infection were the slowest to succumb to infection, averaging 131.1 hours from exposure to death.

Intriguingly, we also noticed that the number of flies killed by *E. muscae* in our 0 hr L:D condition (18.5%) was much lower than that observed in any other condition (60.9%-78.1%) (**Figure 1A**, **Figure 1 - S1B**). Given that we had collected data for each lighting condition from a different independent experiment, it was possible that this was due to a batch effect. That is, low mortality in the 0 L:D experiment might have been observed due to aberrantly low infection rates during the 0 hr experiment. To address this possibility, we exposed flies to *E. muscae* for all six lighting treatments (0 hr, 24 hr, 48 hr, 72 hr, 96 hr and 168 hr L:D) on the same day, housed flies individually under their respective lighting conditions, and counted the number of flies who had succumbed to infection after seven days. Once again, we saw very low mortality among flies from the 0 hr L:D group (8.7%), compared with the remaining treatments (53.5-65.7%) (**Figure 2A**), consistent with our initial observation.

**Figure 2.**
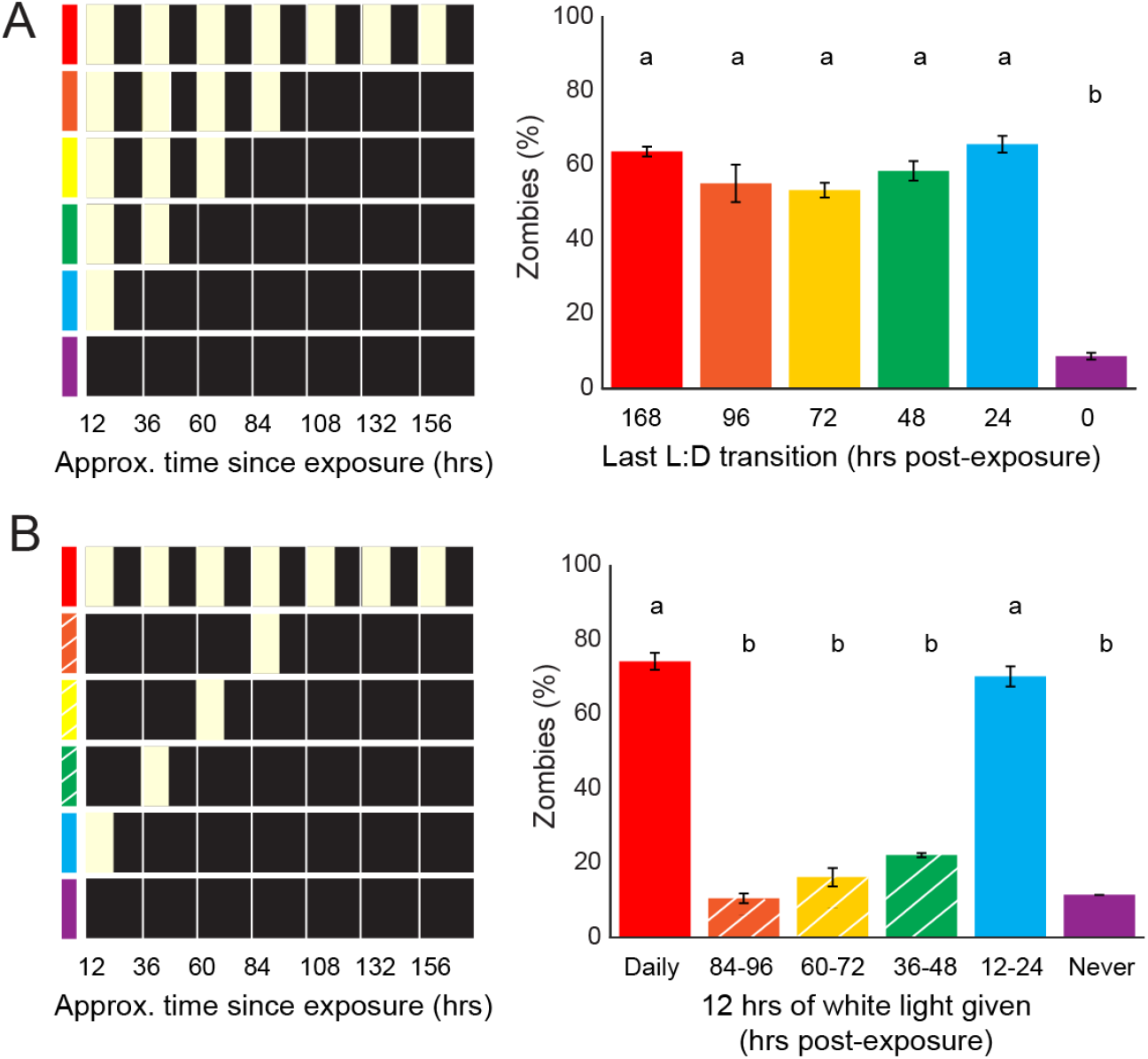
*E. muscae* infection is gated by light cues in the first 24 hours after exposure. A) Right: Percentage of zombies (flies that died and then sporulated) within seven days of *E. muscae* exposure under indicated light conditions at left. B) Right: percentage of zombies subjected to lighting cues diagrammed at left. White hash marks reflect conditions diagrammed at left. Letters above bars indicate distinct significance groups (p<0.05) as determined by pairwise comparisons (two-tailed t-test) for each pair of conditions.

We next wondered if there was a critical period during which light is required for an infection to establish. To test this, we infected flies with *E. muscae* and exposed them to one 12 hour period of light aligned with their entrained photophase within the first 96 hours. Flies that received L:D cues throughout or just in the first 24 hours had high, comparable rates of infection (74.3% and 70.2%, respectively). All remaining groups had low rates of infection (10.5%-22.1%). These data very clearly demonstrated that 24 hours of L:D conditions were necessary and sufficient for high rates of zombification (**Figure 2B**). Altogether, these experiments demonstrate a novel light-dependent critical window for *E. muscae* infection in the first 24 hours of exposure.

### Timing of death by *E. muscae* in D:D does not reflect host genotype

Having established that the timing of death by *E. muscae* is driven internally, we next sought to test the hypothesis that this mechanism is the host circadian clock. Decades of research in fly chronobiology by many research groups has yielded an extensive understanding of the fly clock. Briefly, the core molecular oscillator of *D. melanogaster* circadian clock consists of a transcription/translation feedback loop between the genes *period* (*per*), *timeless* (*tim*), *Clock* (*Clk*), and *Cycle* (*Cyc*) (summarized by ^34^). During the day, Clk and Cyc proteins form a heterodimer that drives the transcription of genes *per* and *tim*. Per and Tim proteins accumulate in the cytoplasm by nightfall, forming heterodimers that are able to translocate to the nucleus. In the nucleus, Per/Tim complexes dissociate leaving Per to bind to *per* and *tim* promoters, blocking transcription driven by Clk/Cyc heterodimers. With the return of daylight, the blue light sensor encoded by *cryptochrome* (*cry*) activates, driving the degradation of tim protein, de-repressing Clk/Cyc transcriptional activation of *per* and *tim* and resetting the clock.

To test the role of the fly clock in driving the timing of death by *E. muscae*, we selected mutants that span circadian phenotypes: two arrhythmic lines (*Clk[jrk]*^35^, *per[01]*^36^) and two genotypes with altered period lengths (*per[S]*^36^ with a 19 hr period, and *per[T610A.S613A]*^37^, with a 30 hr period, hereafter referred to as *per[30]*). Aggregate free running activity patterns for these mutants are shown in **Figure 3A**; results of chi-squared periodogram analysis for all genotypes in this manuscript can be found in **Table S1**.

**Figure 3.**
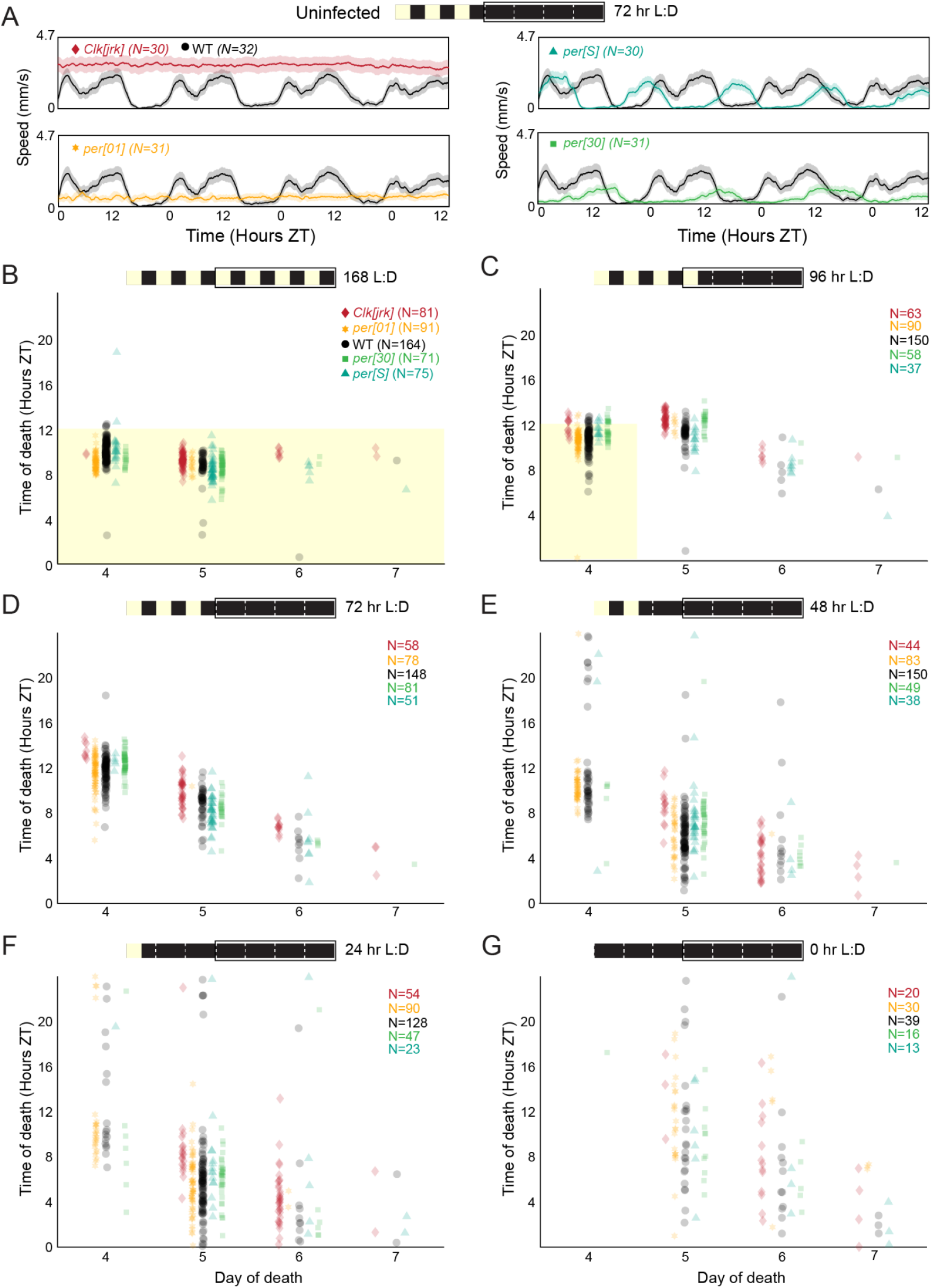
*E. muscae*-infected mutant flies die with similar circadian timing under free-running conditions, regardless of host circadian phenotype. A) Free-running activity patterns of uninfected WT flies and four circadian mutants under D:D conditions. WT = Canton-S, *per[30]* = *per.T610A.S613A*; all other genotypes are named according to their alleles. Flies were raised on a 12:12 white light L:D cycle (ZT0 = 12:00 AM EST) and maintained under this cycle for 72 hours before releasing into D:D and tracking locomotion. B) Time of death for individual flies of four circadian mutants (colors) and WT (black) when kept under L:D conditions throughout infection and tracking. Each point represents the death of a single fly. Each genotype by lighting condition experiment was run once across a six week series of experiments with the exception of WT, which was run twice, once each for two six week series of experiments. C-G) As in B, but flies were released into free running D:D conditions after C) 96, D) 72, E) 48, F) 24, or G) 0 hours after exposure to *E. muscae*. WT data are the same as those presented in Figure 1. For all panels, the schematic at top shows lighting conditions since exposure (time 0, at left), with black box indicating when fly behavior was tracked and corresponding to the plot’s x-axis.

We subjected each of these mutant lines to *E. muscae* exposure and the same set of lighting conditions that we had previously used to test our WT flies (**Figure 1A)**. We observed that the mutants continued to die in a periodic fashion in the absence of light cues, and this timing was consistent across genotypes (**Figure 3B-G**). As with our WT flies, we observed that mutants released into constant darkness 72 hours after exposure or sooner died with periodicity of 20.6 hours (**Figure 3-S1A**), whereas flies exposed to light cues for 96 hours or throughout the experiment showed a mean periodicity of 23.7 hours. As we had also seen before, flies exposed and maintained in the dark throughout infection had much higher survival than flies under any other lighting condition (**Figure 3-S1B**), reiterating the critical role of light during the first subjective day post-exposure in promoting fungal infection. Altogether, these data are consistent with the fly’s molecular clock not dictating the timing of death by *E. muscae*.

To further evaluate the role of the fly circadian system in setting the timing of zombie death, we turned to the unique properties of mutants of cryptochrome, the blue light sensor that conveys light information to the molecular oscillator. For many organisms, including *D. melanogaster*, prolonged housing in constant light (L:L) results in a loss of rhythmicity^38,39^, a consequence of continuous clock-resetting by environmental light. However, *cry[b]* flies have been reported to maintain rhythmicity in L:L conditions^40,41^. We reasoned that, if the fly’s clock was important for determining timing of death, infected *cry[b]* flies would continue to die in a rhythmic manner under L:L conditions while WT flies would die randomly. To test this, we entrained WT and *cry[b]* flies to the same 12:12 L:D cycle, then exposed them to the fungus. After one day in L:D, we transferred them to L:L conditions for the remainder of the experiment. As a control, we ran unexposed flies of each genotype to confirm the expected rhythmic patterns of locomotion (rhythmic for *cry[b]*, arrhythmic for WT). As expected, we observed weakly rhythmic behavior for *cry[b]* flies and clear arrhythmic behavior from healthy WT flies. In contrast, both *cry[b]* and WT flies died with similar timing, with deaths occurring at low levels throughout the day (**Figure 4A)**. This result indicated that the time-keeper for the zombie fly system follows Aschoff’s Rule and loses rhythms in constant light.

**Figure 4.**
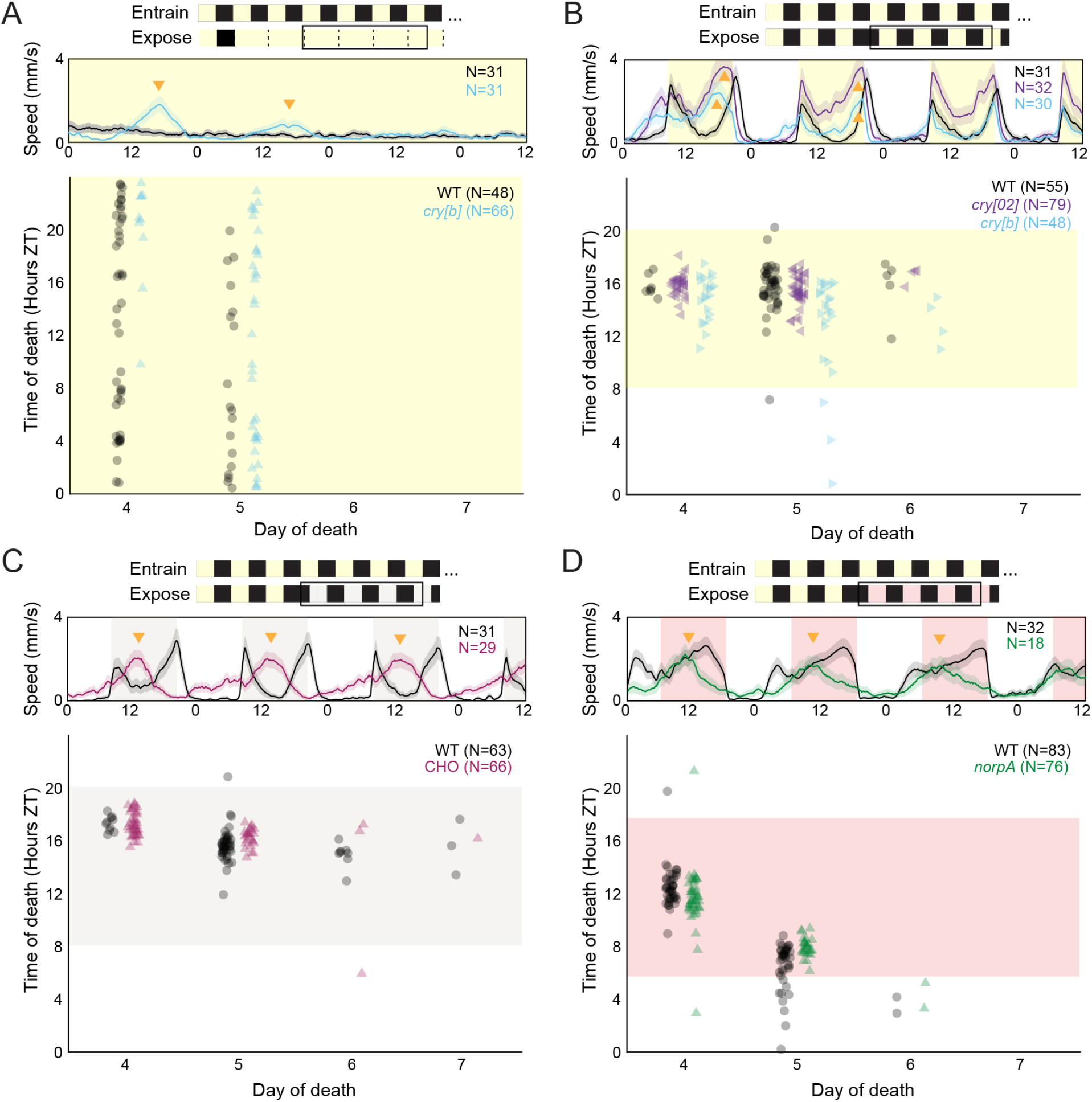
*E. muscae*-infected flies die with similar timing under all lighting conditions tested, regardless of host genetic background. For all panels, the schematic at top summarizes entrainment and experimental conditions. The black outline indicates the period of time during which flies were tracked (corresponding to the middle and bottom panels below). Middle plot shows free-running activity patterns of uninfected flies during the 84 hour window outlined in black. Bottom plot shows time of death for *E. muscae*-exposed flies under these experimental lighting conditions. All flies were entrained on the same 12:12 white light L:D cycle (ZT0 = 12:00 AM EST) prior to experiment. Conditions and genotypes are as follows: **A)** WT (Canton-S) and *cry[b]* flies were exposed to *E. muscae* and, after one day in 12:12 L:D conditions, were housed under constant light (L:L); **B)** WT, *cry[b]* and *cry[02]* flies were exposed to *E. muscae* under 12:12 L:D cues then phase delayed by eight hours on the fourth day after exposure; **C)** WT and *CHO* flies were exposed to *E. muscae* under 12:12 L:D cues then phase delayed with dim white light by eight hours on the fourth day after exposure; **D)** WT and *norpA* flies were exposed to *E. muscae* under 12:12 L:D cues then phase delayed with dim red light by six hours on the fourth day after exposure. Each dot reflects the time of death for an individual fly. Each experiment was run at least twice, with one representative experiment shown above. Yellow shading indicates presence of bright (>1000 lux) white light; tan, dim (40-50 lux) white light; red, dim (<10 lux) red light. Activity data are the mean (solid lines) +/- SEM (shaded regions). Golden arrowheads highlight deviations in activity of uninfected mutant flies relative to uninfected WT flies.

In addition to maintaining rhythms in L:L, *cry* mutants are slow to adapt to light-driven phase delays in their cycle. Whereas WT flies rapidly adjust to a phase delay, *cry* mutants take around three days to realign their behavioral patterns with their new light cues^42^. To test the role of host machinery in dictating timing of death, we exposed WT flies as well as hypomorphic *cry[b]* and null *cry[02]* mutants to *E. muscae* and maintained them on their entrained 12:12 L:D cycle for the first 72 hours after exposure. On the fourth day after exposure, we challenged these flies by delaying the onset of light (i.e., sunrise) by eight hours, resulting in an eight hour phase delay. As has been previously reported, our uninfected WT flies immediately adjusted to these new cues (**Figure 4B**). In contrast, both *cry* mutants showed strong morning anticipation to their previously entrained light cycle on the day of the phase shift as well as premature evening locomotor peaks for the following 48 hours. These activity differences were not reflected in the timing of death across genotypes: all flies died with an approximate eight hour phase delay relative to their entrainment light cycle (∼ZT16). Again, this result is consistent with the host clock not determining the timing of death and suggests that the mechanism driving the timing of death can detect white light.

In all of these experiments, the fly could still sense light. Thus, it was still possible that even if the fly’s clock were not driving the timing of death, that light sensation by the host was involved in setting this timing. To address this, we leveraged flies triply mutant for *cry* and two histamine receptors (*HisCl1* and *Ort*), the latter two genes being responsible for mediating cry-independent photoreception^43^. These “CHO” mutant flies are essentially light insensitive: they cannot see or entrain to low levels of white light. However, they have been shown to entrain to very intense white light, through as yet undetermined mechanisms^43^. We leveraged these features to first entrain CHO flies to a 12:12 L:D cycle under bright white light (>1000 lux). We then exposed WT and CHO flies to *E. muscae*. As with the previous phase shift experiment, we kept exposed flies on their entrainment light cycle for 72 hours before challenging them with an eight hour phase shift delay, this time with dim white light (40-50 lux). As expected, WT flies immediately adjusted to the phase shift, whereas CHO flies continued to follow their previous entrainment, with the evening peak becoming dominant, as is typical for flies kept in free-running conditions^44^ (**Figure 4C**). Despite this, all flies, regardless of genotype, died with similar timing from *E. muscae*, following the phase-shifted light cycle to die at what was previously ∼ZT16. Consistent with our previous experiments, these data support that the time-keeper in the zombie fly system is fly-independent. They also demonstrate that the time-keeper can detect and resynchronize to low levels of white light.

We next wondered if the timekeeper could detect red light. While WT *D. melanogaster* can’t see low-intensity red light, they can use red light cues for entrainment^45^. Mutating the gene *no receptor potential A* (*norpA*), which encodes phosphoinositide phospholipase C, renders flies unable to entrain to red light^46^. To ask if our timekeeper could detect red light, we used WT and *norpA* mutants in a phase-shift challenge experiment utilizing red light. Similar to our previous experiments, we exposed WT and *norpA* mutants to *E. muscae* and maintained them on their entrained 12:12 L:D cycle for the first 72 hours after exposure. We then challenged them with a six hour delayed phase shift, this time with red light (<10 lux). As expected, uninfected WT flies were able to follow the new red light cues, but uninfected *norpA* mutants were not. However, despite these differences in the timing of activity, infected flies of both genotypes died with similar timing, which reflected the original white-light entrainment regime (**Figure 4D**). These data were consistent with what we had previously seen when switching flies housed on 12:12 L:D into D:D after 72 hours (**Figure 3D**) and showed an interdeath interval of ∼20 hours (**Figure 4-S1**). In addition to being consistent with the hypothesis that timing of death is independent of fly genotype, this experiment also shows that the timekeeper in this system cannot see red light.

Altogether, these data strongly suggest that fly circadian and photoreception machinery does not determine the timing of death by *E. muscae*. Additionally, these experiments revealed key attributes of the mechanism responsible for this timekeeping: it maintains a free-running period of less than 24 hours, becomes arrhythmic under constant light, can sense and rapidly adjust to bright or dim white light, and cannot detect red light. Thus, we were driven to consider an alternative hypothesis: could the time-keeper belong to *E. muscae*?

### *E. muscae* shows rhythmic mRNA expression levels in the absence of environmental cues

As a first step to address the hypothesis that *E. muscae* determines the timing of death, we sought to determine if *E. muscae* is capable of endogenously sustaining its own rhythms and not just following host cues (which may serve as Zeitgebers). To do this, we leveraged the unique ability to grow *in vitro E. muscae* cultures in supplemented insect cell medium (Grace’s insect medium supplemented with yeastolate, L-glutamine, lactalbumin hydrolysate, and fetal bovine serum). This medium approximates fly hemolymph but is free of any fly-derived components. Importantly, this setup is entirely artificial - in nature, *E. muscae* does not replicate outside of its host.

Whereas *E. muscae* overtakes the fly host in as few as four days under laboratory conditions, *E. muscae* grows slowly *in vitro*. By spectrophotometric estimation, the doubling time of *in vitro* grown *E. muscae* cultures is 30 hours at room temperature (21°C) (**Figure 5-S1**). Based on these growth kinetics, we subjected *E. muscae* cells over the course of three weeks to a series of lighting conditions to obtain a log-phase culture entrained to a 12:12 L:D cycle (see Methods for details). We then split our log-phase culture and subjected each half to a different lighting condition: either continued incubation with L:D cues or release into D:D. After letting the cells rest for ∼21 hours under their respective conditions, we sampled these cultures every four hours (N = 4 samples per condition per time point) over the course of a 24 hour period, taking great care to avoid introducing light to D:D samples (see Methods). We then extracted and sequenced mRNA, quantified gene-level expression, and used MetaCycle^47^ to identify genes with a cyclic pattern of expression shared between L:D and D:D (**Supplementary File 1**).

**Figure 5.**
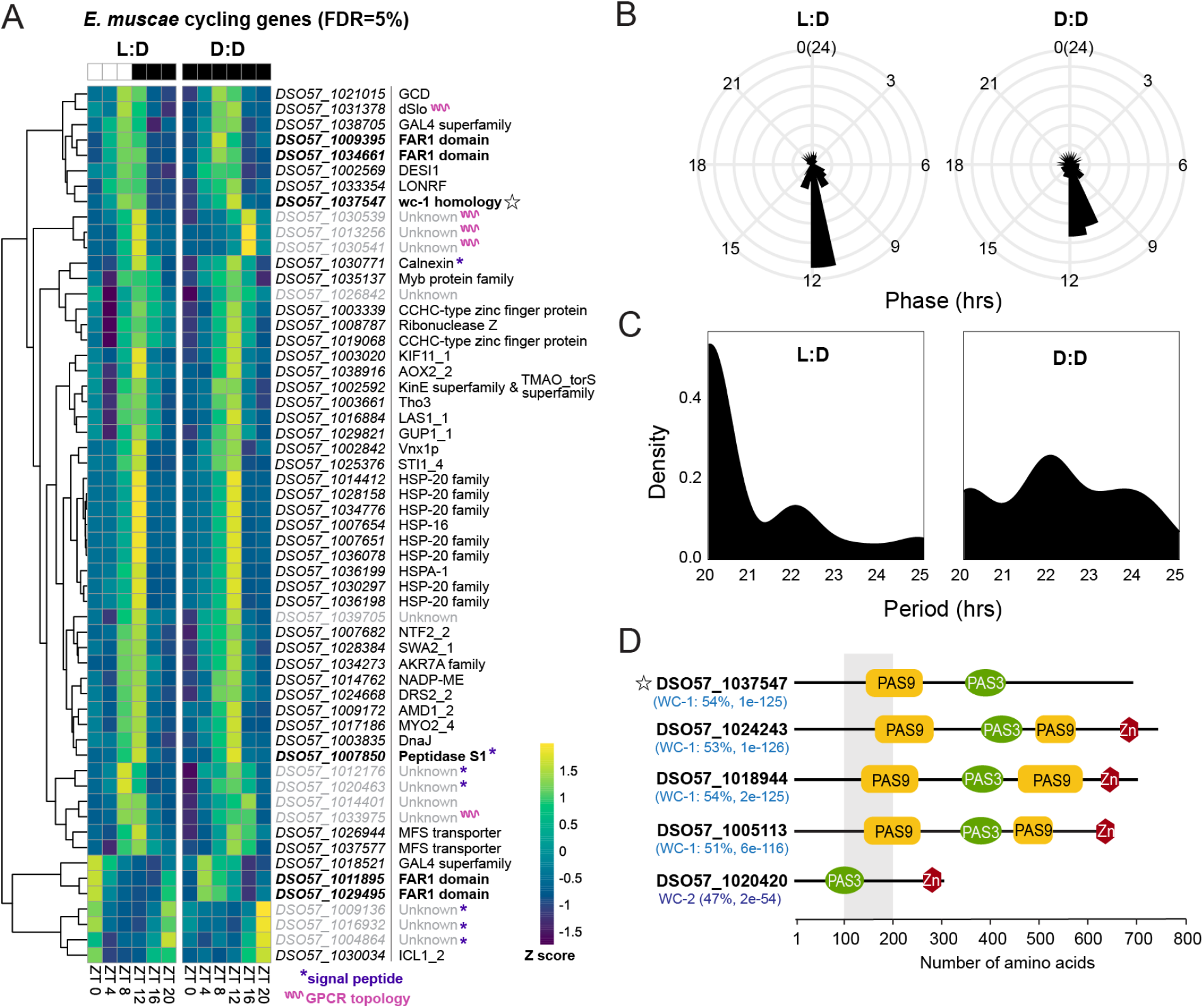
*In vitro E. muscae* shows rhythmic gene expression under both L:D and D:D conditions. A) Gene expression heatmap of Z-scored rhythmic transcripts from *in vitro*-grown *E. muscae* that cycle under both L:D and D:D conditions (MetaCycle, false discovery rate (FDR) = 5%, minimum period = 20 hours). Genes are clustered according to expression patterns. Scale bar for Z-scores given at bottom right. Star = putative *wc-1* homolog; purple asterisks = gene has predicted signal peptide, pink squiggle = gene product has predicted G-protein coupled receptor seven transmembrane (GPCR 7TM) topology. B) Phase and C) period distribution of cycling genes shown in A. D) Predicted *E. muscae* proteins with high homology to *Neurospora crassa* White Collar 1 (WC-1) and White Collar 2 (WC-2), as determined by BLASTp (query coverage 79%-87%). Multiple isoforms were identified for each protein; domain architecture of the longest isoform is shown. Percent positive matches to *N. crassa* homologs given in parenthesis, along with BLASTp E-values. PFAM domain architecture is shown to the right. Star denotes putative *wc-1* homolog observed to cycle under both L:D and D:D conditions at FDR 5 (same as panel A).

We detected 57 genes that cycled in both L:D and D:D in our *in vitro* cultures at a false discovery rate of five percent (**Figure 5A, Supplementary File 1**). In both conditions, the majority of cycling transcripts peak in expression at or around ZT12 (subjective sunset and coincident with the peak of deaths) (**Figure 5B**), however there was a notable minority of genes that were phase-advanced in D:D compared to L:D, peaking at ∼ZT11 instead of ZT12. The average period for all cycling genes was <24 hours under both L:D and D:D conditions, with means at 20 and 22 hours, respectively (**Figure 5C**).

We next examined our cyclers to look for common functional patterns. Based on Pfam, InterPro, EggNog, and GO annotations, we grouped these genes into seven groups: chaperones (14 genes), DNA binding (9 genes), metabolism (9 genes), transport (7 genes), RNA processing (3 genes), signaling (3 genes) and unknown (12 genes). These genes and their phases are summarized in **Table S2**.

Notably, two genes (*DSO57_1031378* and *DSO57_1037547*) within the signaling group contained PAS domains. In performing a BLASTp search on *DSO57_1037547,* we were surprised to find the predicted protein had strong homology to *wc-1* (**Figure 5D**). This result was unexpected because our previous domain-level analysis of a panel of entomophthoralean genomes had identified only one putative homolog of *white collar 1* (*wc-1*) in *E. muscae* (*DS057_1005113*)^48^.

In addition to the *wc-1* homolog that we uncovered in our list of cyclers, we found two additional putative homologs of *wc-1* – *DSO57_1024243* and *DSO57_1018944* – for a total of four *wc-1* homologs (**Figure 5D**). Among these four homologs, *DSO57_1037547* was the only gene we observed cycling below an FDR cut-off of five percent, however *DSO57_1005113* and *DSO57_1024243* were detected as cyclers when relaxing the FDR to 10% (**Figure 5-S2**). *DSO57_1018944* is expressed at very low levels *in vitro* and was not observed to cycle in L:D or D:D. Each of these putative *wc-1* homologs includes not one, but two PAS_9 domains in addition to one PAS_3 and one GATA domain. Our reanalysis confirmed a previously-identified homolog candidate, *DS057_1005113*^48^. We also discovered a putative *wc-2* homolog, *DSO57_1020420*, which was not observed to cycle across either L:D or D:D condition (**Figure 5-S2**).

Overall, our transcriptomic data support the conclusion *E. muscae* can endogenously keep time with a free-running phase of ∼22 hours and, *in vitro,* most cycling genes are phased for peak gene expression at sunset, coincident with the stereotyped timing of host behavior manipulation and death. Combined with our fly behavioral data, these findings suggest that *E. muscae* possess the circadian clock that dictates when its host dies.

## Discussion

For decades, it has been known that *E. muscae* only kills at sunset^32^, but the basis for this eerie punctuality has remained undetermined. Leveraging the experimentally tractable zombie fruit fly system, we provide the first mechanistic insights as to how the specific timing of death is achieved. Our findings are consistent with the hypothesis that *E. muscae*, not the host, drives this timing via an endogenous fungal clock.

### Death by *E. muscae* follows an internal timekeeper

By tracking fly locomotion under a range of L:D conditions, we observed that death of WT flies infected by *E. muscae* continued to occur in a time-restricted fashion in the absence of light cues (**Figure 1A, B**). We observed an average free-running period of death (the time elapsed between median death on consecutive days) of 20.8 hours (**Figure 1C**). In our hands, WT flies have a typical free-running period of ∼23.5-24 hours (**Table 2**); the observed free-running periodicity is already inconsistent with that maintained by the fly. These data reveal that there is a mechanism intrinsic to the infected fly system that drives the timing of death, and hint that this mechanism is fly-independent. Notably, Krasnoff and colleagues were the first to observe that death under constant darkness (D:D) still occurred with a roughly daily rhythm in house flies, suggesting the involvement of a biological clock^32^.

Circadian systems are defined by three key features: a) a free-running period of approximately 24 hours, b) persistence in the absence of environmental cues, and c) stability across biologically relevant temperatures-a phenomenon known as “temperature compensation”^39^. While our data fulfill the first two criteria, we have not yet addressed whether *E. muscae*’s rhythms are temperature-compensated. We have previously found that both summiting behavior and fungal outgrowth are strongly inhibited by modest temperature increases (28°C)^31^. To enable rigorous testing of temperature compensation, a more refined delineation of the temperature range that supports normal fungal growth and host manipulation is needed.

### *E. muscae* infection requires light

Initially, we had sought to test whether environmental light cues are required for exposed flies to time their death by comparing infections under L:D and D:D conditions. However, we encountered a complication: infection rates were extremely poor when flies were housed in constant darkness (**Figure 1-S1**). This unexpected result led to the discovery that light is needed during *E. muscae* exposure for successful infection (**Figure 2A**). Further work pinpointed this critical light-sensitive window to the first 24 hours following exposure (**Figure 2B**).

We have shown that 12 hours of white light during the first day post-exposure to *E. muscae* is sufficient to enable infection, but it remains unclear whether this period of light exposure can be shortened with the same effect. We also have yet to explore if specific wavelengths alone (e.g., just blue light) can enable infection. In addition, it remains unclear why light is required in order for infection to establish and whether the requirement for light is on the side of the fungus, fly, or both. While it’s formally possible that failure for infection to take hold in darkness could be due to fly machinery (for example, via an increased immune response in darkness vs. light), we favor the hypothesis that the need for light rests on the side of the fungus. This thinking is motivated by published reports describing *E. muscae*’s photosensitivity^49^, the presence of several light-sensitive domains in the *E. muscae* genome^48^, and unpublished observations in our lab (J. Tabin, C. Eaton, C. Song personal communication). Future studies are needed to address why and how light plays such a critical role in establishing *E. muscae* infection.

### Timing of fly death is not set by the host

Following the observation that death remains periodic in the absence of light cues, we leveraged the power of the fruit fly genetic toolkit to test the role of the fly’s clock in setting the time of death. We observed that fly mutants spanning a range of circadian and vision phenotypes died with similar timing as WT controls across light conditions (**Figure 3B-G**, **Figure 4A-B**). That is, we did not observe differences in the timing of fly death that corresponded with host genotypic effects on the host’s circadian rhythm. This suggests that fly pathways do not impact *E. muscae* induced death. Our finding stands in contrast to the existing literature on parasite-mediated behavioral manipulation, where the behavioral shifts have either been shown or hypothesized to run through, hijack, or otherwise rely upon the host’s intact circadian machinery (e.g.,^22,28,30^). In this system, *E. muscae* appears to dispense with the host clock altogether, establishing its own independent axis of timekeeping.

### The timekeeper can sense white, but not red, light

We next addressed the role of fly photoreception in timing of death. First, we compared the timing of death of two photoreception mutants (CHO and *norpA*) to WT and, again, failed to see that death timing was affected by fly genotype. CHO flies, triply mutant for *cry, Ort*, and *HisCl*, are unable to entrain to dim white light^43^. When CHO flies and WT flies were challenged with an eight hour phase delay using dim white light, fly activity was clearly different between mutant and WT flies but timing of death was similar between genotypes (**Figure 4C**). This result demonstrates that the timekeeper in the zombie fly system can detect and entrain to white light and supports our conclusion that the host does not drive the timing of death.

Similarly, we compared the timing of death of *norpA* flies (unable to entrain to red light) to that of WT flies (able to entrain to red light). Flies of both genotypes die with similar timing, despite clear differences in the locomotor activity of uninfected animals under these conditions (**Figure 4D**). This result again supports the conclusion that host phototransduction does not influence when death occurs. Importantly, the periodicity of death in this experiment approximated what we observe under D:D conditions (**Figure 3D-F**), indicating that the timekeeper cannot detect red light.

Intriguingly, the predicted photosensitivity of *E. muscae* – capable of detecting blue, but not red light – aligns with the empirically-determined properties of the timekeeper. Domain-level analysis of the *E. muscae* genome predicts that the fungus can detect blue wavelengths (likely mediated by *wc-1* as well as genes encoding FAD binding 7 and DNA photolyase domains), but lacks any phytochromes, which detect red light^48^. Although these genomic features do not offer conclusive proof that *E. muscae* keeps time, they provide a strong mechanistic rationale for our observations and heavily favor a model where *E. muscae* directly senses light cues to time host death.

### *E. muscae* shows endogenous gene expression rhythms

To determine if *E. muscae* can maintain ∼24 hour rhythms in the absence of zeitgebers, we measured transcripts from *in vitro* fungal cultures every four hours over the course of a day under both L:D and D:D conditions. We observed 57 genes that exhibit statistically significant cycling across both conditions (**Figure 5A**). These genes showed similar phases (**Figure 5B**) and periods (**Figure 5C**) across both L:D and D:D conditions, supporting the interpretation that *E. muscae* can maintain endogenous rhythms. Other studies have observed a similar number of transcripts that cycle across experimental conditions^50,51^. Thus, our results are well-aligned with other studies on fungal rhythms.

Importantly, *E. muscae* is an obligate pathogen that is never found naturally growing outside of the fly host. While we can culture *E. muscae* in the absence of a host in the lab, *in vitro* grown fungal cells likely differ from their *in insecta* grown counterparts. Indeed, whereas *E. muscae* infects with a handful of spores that can completely take over the fly hemocoel within a period of 4-5 days, *in vitro* cultures grow very slowly, doubling only every ∼30 hours (**Figure 5-S1**). The large disparity in division times indicates that our *in vitro* condition was not optimized for *E. muscae* growth. In addition, unlike cells grown inside the fly, our *E. muscae in vitro* cultures are extraordinarily heterogeneous. While all cells are protoplastic, their individual morphologies vary widely within a culture^52,53^. We suspect this variability arises from a lack of host cues helping to coordinate cell physiology as well as a lack of physical confinement – a 20 mL tissue culture flask presents orders of magnitude more volume than the interior of a fruit fly. We hypothesize that this morphological heterogeneity is underpinned by heterogeneity in gene expression. If so, we might expect that all but the strongest periodic signals to be undetectable in circadian transcriptomic data. Future work looking at transcripts that cycle over the course of infection will be needed to characterize the complete range of daily oscillations in *E. muscae* in its natural environment.

Within the genes we observed to cycle *in vitro*, putative functions could be assigned to 45 of 57 transcripts based on conserved domain structures and/or amino acid sequence homology (**Table S2**). Among these, dSlo (*DSO57_1031378*) caught our attention. In *Drosophila*, dSlo or Slowpoke is a BK calcium channel that functions in maintaining circadian output rhythms^54^. Slowpoke activity is in turn modified by the Slowpoke binding protein, Slob^55^. *slob* transcripts have been shown to cycle ^56^. Similarly, work in mice has also demonstrated that the BK calcium channel modulates spontaneous firing rates in a circadian manner^57^. Seeing this annotation for a cycling transcript, we are curious if dSlo may be involved in regulating fungal circadian rhythms. As BK calcium channels have not yet been reported in Dikarya^58^, the putative functions of this gene remain unexplored.

ICL1_2 (*DSO57_1030034*), a predicted isocitrate lyase, also stood out to us owing to its disparate phasing compared to the majority of other transcripts. ICL catalyzes the first step in the glyoxylate cycle, cleaving isocitrate into glyoxylate and succinate. In bypassing the TCA cycle, it allows organisms to avoid losing carbons as CO_2_ and use two carbon precursors for gluconeogenesis and other biosynthetic pathways. In addition to its known role in metabolism, ICL has been shown to play a role in fungal virulence across several species^59,60^. We wonder if cyclic ICL expression could mediate daily fluctuations in virulence and perhaps help *E. muscae* align periods of highest virulence to correspond with entry of the host.

Twelve cycling transcripts lack predicted functions, though five are predicted to be secreted. Three of these are shorter than 300 amino acids—*DSO57_1009136* (153 aa), *DSO57_1004864* (194 aa), and *DSO57_1016932* (232 aa)—making them candidates for small secreted proteins (SSPs). In other fungal pathogens, SSPs often act as effectors that significantly influence host physiology^61^, so it is tempting to speculate that molecules released by *E. muscae* may mediate fungal-fly interactions. Curiously, all three of these genes were antiphase to the majority of cycling genes, peaking around ZT0 rather than ZT12. Future experiments measuring *in insecta* fungal gene expression are a much needed first step towards addressing this hypothesis.

### Insights into the posited *E. muscae* clock

This study led us to discover three additional putative *wc-1* homologs (*DSO57_1024243*, *DSO57_1018944*, and *DSO57_1037547*) as well as a novel putative *wc-2* homolog (*DSO57_1020420*) (**Figure 5D**), which had been overlooked in a previous domain-based genomic analysis^48^. We suspect that these homologs were previously missed due to a versioning difference in PAS annotation (e.g., the domain was annotated as PAS rather than PAS_3 specifically). The presence of multiple *wc-1* homologs may enable *E. muscae* to mount specific light responses to different light cues, as is the case for mucoromycete fungi *Phycomyces blakesleeanus* and *Mucor circinelloides*^62,63^. Indeed, many non-dikarya fungi possess multiple copies of *wc-1.* These are thought to have arisen from gene duplications and are speculated to enable specialized responses to different wavelengths and intensity of light^64^. Future functional studies will be critical in defining roles for these different homologs over the *E. muscae* life cycle.

Among our newly identified *wc-1* homologs, *DSO57_1037547* was the most robust cycler *in vitro*. Intriguingly, *DSO57_1037547* lacks a GATA zinc finger domain, suggesting that the resultant protein cannot directly bind DNA to drive transcription of clock-controlled genes and is unlikely to function precisely as *wc-1* does in *N. crassa*. Given that *DSO57_1037547* retains a PAS_9 domain, we expect its gene product can form protein-protein interactions. Indeed, several basidiomycete species including *Pleurotus ostreatus*, *Schizophyllum commune*, and *Cryptococcus neoformans*, encode *wc-1* homologs that lack a GATA domain, all of which have been implicated in fungal development and been shown to physically interact with their predicted WC-2 proteins^65–67^. We suspect that *DSO57_1037547*’s gene product may function similarly in *E. muscae*.

Rhythmic gene expression *in vitro* along with the expression of putative homologs of *wc-1* and *wc-2* are strong reasons to suspect that *E. muscae* has a circadian clock. Interestingly, *E. muscae* lacks obvious candidates for homologs of *frequency* (*frq*) and *frequency-interacting RNA helicase* (*frh*), which comprise the negative arm of the *N. crassa* oscillator^48^. This raises intriguing questions about how a potential molecular oscillator could be structured in *E. muscae*. *E. muscae* is not alone in this regard: several fungi with circadian phenotypes under free-running conditions lack a *frq* homolog. These include the ascomycetes *Aspergillus flavus* and *Cercospora kikuchii*, the basidiomycete *Neonothopanus gardneri*, and the mucoromycete *Pilobolus sphaerosporus*^68,69^. Thus, it is clear that there must be *frq*-independent oscillators within fungal lineages, and the players remain open for discovery. We think *E. muscae* provides an exciting platform to identify these new mechanisms.

The prevailing view is that the negative arm of all fungal and animal clocks are phosphoproteins that function similarly to *frq* and *per*^70^. None of the genes within our list of *in vitro* cyclers seemed a likely candidate for *frq*, and frustratingly, the defining properties of *frq —* intrinsic disorder and lack of sequence conservation — preclude us from proposing any possible candidates in *E. muscae* based solely on the annotated genome^71^. We expect that measuring gene expression over circadian time *in insecta* will be critical in revealing a broader set of cyclers, which may help us to identify candidates for the negative arm of *E. muscae*’s putative oscillator.

### Death according to a fungal clock? Evolutionary considerations and broader implications

Our data show that 1) flies consistently die at a specific time of day (**Figure 1, 3, 4**), 2) this timing persists in the absence of environmental cues (**Figure 1 & 3**), and 3) this timing is not impacted by disruption of host circadian or photoreception systems (**Figure 3 & 4**). Furthermore, our data demonstrate that *E. muscae* can maintain autonomous transcriptional rhythms (**Figure 5**), consistent with an interpretation that the fungus keeps time. The most parsimonious explanation for these findings is that *E. muscae* sets the timing of host death.

The question arises: why does *E. muscae* kill at sunset? Any potential explanation for this phenomenon needs to take into consideration that though the host dies at sunset, the fungus does not emerge from the host and begin to sporulate until some hours after, with most spores fired within the first 12 hours after death^72^. The timing of fungal emergence and sporulation is likely what death at sunset seeks to optimize. Given how critical the timing of fungal emergence is for fungal fitness, it could follow that the fungus would evolve mechanisms to control it directly rather than relying on a timekeeper within the infected host.

Sporulation at night could favor fungal fitness for many possible reasons, but generally falls into two broad categories: optimization of host interaction and optimization of environmental conditions. As an obligate pathogen, *E. muscae* must find new hosts to survive. One possibility is that *E. muscae* sporulation is timed to coincide with periods of highest infection susceptibility in flies. However, studies looking at susceptibility to bacterial pathogens in flies over the course of a day have found that flies are most resistant to infection after dark^73^. While it is possible that immunity to fungal infections may differ, this seems unlikely to be the primary driver of *E. muscae*’s timing. Alternatively, sporulation at night may be advantageous in targeting sleeping hosts, both because they are immobile and unable to groom. However, *E. muscae* produces not just primary conidia (the first wave of spores that shoot off of dead flies), but regularly forms secondary conidia^74^. It is possible that the formation and firing of secondary and potentially even higher order conidia may coincide with the morning peak of fly activity thereby increasing collision events between spore and fly, though this remains speculative.

*E. muscae*’s timing may also bias spore emergence to occur in favorable environmental conditions. Like many fungi, *E. muscae* thrives in cooler, damp conditions. Indeed, studies measuring sporulation, germination, and host infectivity over a range of temperatures and humidities consistently point to a preferred temperature of ∼21°C with saturating ambient humidity^53^. The longest period of the day during which such conditions occur is after the sun goes down. In addition, air turbulence has been shown to be minimized during the night^75^. This could allow spores to settle within hours, rather than days, thereby maximizing contact of viable spores with suitable hosts. Finally, *E. muscae* is unpigmented so lacks protection against UV irradiation. Recent work on photoinhibition of fungal development supports the idea that circadian scheduling might help fungi avoid damaging light exposure^76^. Given the inconsistencies with the hypotheses around how this timing would favor host interactions and the strong data measuring *E. muscae*’s abiotic tolerances, we tend to favor abiotic pressures as the primary motivation for *E. muscae* to emerge at night.

Whether the host’s own circadian clock is impacted by *E. muscae* remains an open question. Because mortality began within 24 hours of data collection, we could not fully evaluate what becomes of host rhythms in infected individuals. Given the robustness of free-running fly rhythms in uninfected counterparts for our experiments (**Table 2**), it is all the more striking that our data did not show the host clock influencing the time of death. This suggests *E. muscae* might somehow suppress host rhythms, a hypothesis that future studies should evaluate.

While our data consistently point to the host’s clock not being required to determine the timing of death, conclusive evidence that *E. muscae’s* clock drives this timing will require genetic manipulation of the fungus. Indeed, current work in our lab is focused on developing transgenic tools for *E. muscae.* Functional knockouts of the core fungal clock components (such as the *white collar* genes) will provide the ultimate evidence of whether death by *E. muscae* is timed by the fungus.

Nevertheless, fungal genetics are not needed to be confident that *D. melanogaster* does not drive the timing of its death by *E. muscae*. Consequently, death regulated by a fungal, not a fly, clock should be established as the *de facto* null hypothesis in this system. Our work raises broader implications for manipulation of host behavior in *E. muscae*’s closest relatives, obligate insect pathogens of the family Entomophthoraceae. Fungi in this clade infect a variety of insects and have almost uniformly been observed to induce time-of-day-specific behavioral phenotypes and death in their hosts^53,77,78^. Given the near ubiquity of daily timed events in this fungal family and the strong evolutionary pressure to optimize dispersal, we suspect that fungal-driven circadian timing of host behaviors and death is prevalent across the Entomophthoraceae.

In conclusion, our work points to the use of a pathogen’s endogenous clock to schedule host death. The evolutionary rationale is clear: by dictating temporal control of host death, *E. muscae* ensures its transmission strategy unfolds under the most favorable conditions. We anticipate that this kind of pathogen-driven circadian timing may be far more common than currently recognized, especially among behavior-modifying parasites with narrow transmission windows.

## Acknowledgements

We are indebted to Dr. Stan Lazopulo and Dr. Kyobi Skutt-Kakaria for their generous gifts of CHO and *norpA* mutant flies, respectively. We used FlyBase to find information on available circadian mutant lines^79^. Fly lines were obtained from the Bloomington Drosophila Stock Center, which is supported by the NIH (P40 OD018537). We thank the Bauer Core Facility at Harvard University for RNAseq library preparation and sequencing. The sequence-based computations in this paper were run on the FASRC FASSE cluster supported by the FAS Division of Science Research Computing Group at Harvard University. We are grateful to Dr. Kevin Tsui and Dr. Venki Murthy for their helpful comments and suggestions. CNE also extends her gratitude to her long-time collaborator Nora the dog, who was essential in warming her lap while she drafted this manuscript.

## Author CRediT statement

**LTU:** Methodology, Software, Validation, Formal analysis, Data curation, Writing - Review & Editing, Visualization

**DR**: Investigation, Formal analysis

**AM**: Investigation

**FB**: Investigation

**BdB:** Resources, Writing - Review & Editing, Supervision, Funding Acquisition

**CE:** Conceptualization, Methodology, Software, Validation, Formal analysis, Investigation, Resources, Data Curation, Writing - Original Draft, Writing - Review & Editing, Visualization, Supervision, Project Administration, Funding Acquisition.

## Methods

### Fly stocks and husbandry

All fly stocks were maintained in vials on cornmeal-dextrose media (11% dextrose, 3% cornmeal, 2.3% yeast, 0.64% agar, 0.125% tegosept [w/v]) at 21°C and ∼40% humidity in Percival incubators under 12 hours light and 12 hours dark lighting conditions (12:12 L:D; >=500 lux) and kept free of mites. All fly stocks used for experiments are listed in the **Key Resources Table**.

### *E. muscae* husbandry

A continuous *in vivo* culture of *E. muscae* ‘Berkeley’ (referred to herein as *E. muscae*; USDA ARSEF#13514) isolated from wild Drosophilids ^33^ was maintained in Canton-S flies cleared of *Wolbachia* bacteria following the protocol described in Elya et al., 2018 and summarized as follows. Canton-S flies were reared in bottles containing cornmeal-dextrose media (see Fly stocks and husbandry) at 21°C and ∼40% humidity under 12:12 L:D. *E. muscae*-killed flies were collected daily between ZT14 and ZT18 using CO_2_ anesthesia. To infect new Canton-S flies, 30 fresh cadavers were embedded head first in the lid of a 60 mm Petri dish filled with a minimal medium (autoclaved 5% sucrose, 1.5% agar prepared in milliQ-purified deionized water, aka ‘5AS’). Approximately 330 mg of 0–5 day-old Canton-S flies were transferred to a small embryo collection cage (Genesee #59–100, San Diego, CA) which was topped with the dish containing the cadavers. The cage was placed mesh-side down on a grate propped up on the sides (to permit airflow into the cage) within an insect rearing enclosure (Bugdorm #4F3030, InsectaBio, Riverside, CA) and incubated at 21 °C, ∼40% humidity on the same 12:12 L:D cycle as used for all fly rearing. After 24 hr, the cage was inverted and placed food-side down directly on the bottom of the insect enclosure. After 48 hours, the cadaver dish was removed from the cage and replaced with a new dish of 5AS without cadavers. Starting at 96 hr, the collection cage was checked daily for up to four days between ZT14 and ZT18 for E. muscae-killed flies. These were collected using CO_2_ anesthesia and used to infect additional flies for experiments as described below.

### Measuring time of death by *E. muscae*

Experimental flies were exposed to *E. muscae* as follows: eight sporulating Canton-S cadavers were embedded in a 2.3 cm-diameter disc of ∼3.5 mm thick 5AS that was transferred with 6” forceps into the bottom of an empty wide-mouth Drosophila vial (Genesee #32–118). A ruler was used to mark 1.5 cm above the top of the disc. 0–5-day-old flies of the experimental genotype were anesthetized with CO_2_, and 35 (∼half male, ∼half female) were transferred into the vial. The vial was capped with a Droso-Plug (Genesee #59–201) which was pushed down into the vial until the bottom was level with the 1.5 cm mark. For each experimental tray, three vials of flies were prepared in this way to expose a total of 105 flies; one additional vial of 35 flies was prepared identically but omitted cadavers as a non-exposed control. Together, these four vials were sufficient to fill a tray of 128 arenas (see Measuring locomotor activity). All prepared vials were incubated in a humid chamber (a small tupperware lined with deionized water-wetted paper towels) at 21°C. Vials to be exposed to 12:12 L:D white light cues were housed directly in the fly rearing incubator. Vials to be kept in constant darkness (D:D) were transferred to film changing bags (VANZAVANZU, Amazon.com) that were housed in the fly rearing incubator. Flies to experience constant light (L:L) were transferred the second day following exposure to a behavior enclosure with constant illumination (∼150 Lux) from 5300K white LEDs.

After 24 hr, the vials were removed from the humid chamber, and the Droso-plugs were pulled to the top of the vial to reduce fly crowding. For vials kept in D:D, flug raising was done under dim red light (<20 lux).

Under 12:12 L:D cues at 21°C, flies first die from *E. muscae* infection starting ∼5 hours prior to sunset on the fourth day after exposure. Flies continue to die until seven days after exposure, always at sunset, though the majority of deaths occur on days four and five. To capture all possible deaths due to *E. muscae*, flies were anesthetized with CO_2_ after 72 hours and loaded individually into behavior arenas prepared with 5AS as described in ^31^. For experiments in which environmental light cues were still being provided at 72 hours, flies were loaded into behavioral arenas during photophase (the light period of their entrained 12:12 L:D circadian cycle) under ambient light. Flies that were not to experience white light cues were loaded into chambers under red light. Using red light, behavior trays were placed in summit assay boxes and flies were tracked starting between ZT17 and ZT20. Tracking proceeded for ∼96 hours (through the seventh day following exposure), ending on ZT13 four days from the start of tracking.

Tracking data were collected at 3 Hz using MARGO v1.03 ^80^; https://github.com/de-Bivort-Lab/margo) using infrared illumination with the following settings: tracking threshold = 18, minimum area = 10, min trace duration = 6. Default settings were used for other configuration parameters. Environmental lighting cues (white or red light) were controlled through MARGO with the help of Chrome Remote Desktop (Google). Unless otherwise stated, white light cues were provided at >=500 lux and red light was provided at <10 lux. Lighting exposure was verified for each experiment via webcam recording of the room in which experiments took place. After tracking concluded, flies were manually scored as either alive (coded as survival = 1 and outcome = 0), dead with evidence of *E. muscae* sporulation (survival = 0, outcome = 1), or dead with no *E. muscae* sporulation (survival = 0, outcome = 0). These annotations were saved in a metadata spreadsheet accompanying each MARGO output file and used in downstream analyses. Only flies that died with evidence of sporulation were considered “zombies” and used for analysis. Time of death was called manually for every zombie using behavioral tracking data by CE as described in ^31^. Times of death were scored blind to the experimental group.

Activity data from uninfected flies was used to confirm circadian entrainment and expected response across genotypes to the given experimental lighting conditions. For varied L:D experiments (**Figure 1** and **Figure 3**) each genotype by lighting condition combination was run once (N=96 flies exposed to *E. muscae*, N=32 uninfected flies), except for Cs experiments, each of which were run twice. For each panel in **Figure 4**, at least groups of flies (N=96 flies exposed to *E. muscae*, N=32 uninfected flies) were assayed during separate experiments on separate days and gave similar results; data from one representative experiment is shown. Note that while, conventionally, ZT becomes circadian time (CT) in free-running conditions, we have used ZT throughout the paper to keep axis labels consistent; the presence of lighting cues is indicated by shading.

### Measurement of light intensity

Light intensity was measured using the Lux Light Meter and Arduino Science Journal apps on Pixel 4a and 9 phones as well as using SparkFun OPT4048 Light Sensors (SparkFun, SEN-22638). Readings were consistent across all measurement paradigms.

Rhythmicity analysis of uninfected flies

Rhythmicity of all individual control flies was determined by chi-squared analysis, binning speed data into 60-minute intervals and evaluating period lengths ranging from 18 to 30 hours. Missing data points (NaNs) were replaced with zeros prior to analysis. Speed data were downsampled by summing values within each 60-minute bin to generate a condensed time series of length *N*.

The chi-squared statistic (***Q_p_***) was calculated for each test period (***P***, in 0.1-hour increments) to determine the strength of the rhythmicity relative to total variance, using the following formula:

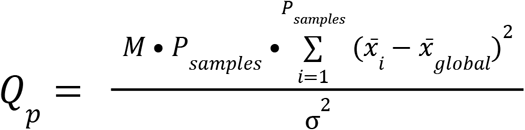

where ***P_samples_***represents the number of bins spanning a given period, ***M*** is the number of full periods that fit within the truncated time series, ***x _i_***is the mean value of the i-th time point across all cycles, ***x _global_***is the global dataset mean, and ***σ***^2^ is the total variance of the dataset.

To establish a conservative significance threshold and correct for multiple comparisons across the tested period range, a permutation test was performed. For each individual fly, the binned time series was randomly shuffled 1,000 times to disrupt any underlying circadian structure while preserving the underlying distribution of values. A chi-squared periodogram was computed for each randomized iteration. The empirical significance threshold (α = 0.05) across the period range was defined as the 95th percentile of the maximum ***Q_p_***values obtained across the permutation matrix, adjusted for the total number of data blocks. A fly was deemed significantly rhythmic if its true peak ***Q_p_*** value exceeded this empirically derived threshold.

### *In vitro* growth of *E. muscae*

Spores from single *E. muscae*-killed Canton-S sporulating cadavers were collected in 60 mm sterile petri dishes using the ascending sporulation method ^81^. Spores were collected in supplemented Grace’s Insect Media (ThermoFisher Scientific #11605094) with added 5% fetal bovine serum (ThermoFisher Scientific #10437010) and grown in volumes of 20 mL in a 25 cm^2^ untreated, vented TC flasks (hereafter: TC flasks; Corning #353014) at room temperature (∼21°C). Cultures were grown without shaking, with flasks laid horizontal to maximize surface oxygen exchange.

### Measuring doubling time of *in vitro* grown *E. muscae*

Three 175 cm^2^ TC flasks containing 150 mL of supplemented Grace’s media with 5% FBS were each inoculated with 3.5 mL of log-phase *E. muscae* culture. Once daily for eleven consecutive days, cultures were gently agitated by up-and-down pipetting with a serological pipette before removing 4.5 mL to sample culture growth. Each 4.5 mL sample was split into 1.5 mL aliquots, dispensed into 1.7 Eppendorf tubes, and pelleted by centrifugation at room temperature for 5 minutes at maximum speed (12,800g). After removing the supernatant was removed by micropipette, the cell pellet was resuspended in 1 mL 1x PBS, resulting in a crude *E. muscae* cell lysate. Lysate was transferred to a plastic cuvette and OD600 was measured (Implen Diluphotometer).

### Measurement of *in vitro* fungal gene expression

In preparation for circadian sampling of *in vitro* grown cells in each L:D and D:D conditions, a new log-phase culture (Culture 0) was used at 1:40 dilution to inoculate a fresh 20 mL culture (Culture 1) in a 25 cm^2^ TC flask. Culture 1 was subsequently incubated at 21°C under constant light for seven days to drive arrhythmia across the culture. Culture 1 was then used at 1:40 dilution to inoculate a fresh 640 mL culture (Culture 2). Immediately after inoculation, Culture 2 was subdivided into four 160 mL cultures each housed in 175 cm^2^ TC flasks and grown under 12:12 L:D at 21°C for seven days. On the seventh day at ZT15, the four flasks containing Culture 2 were gently recombined and mixed. Culture 2 was then split into 30 aliquots of 20 mL and each aliquot was transferred to a fresh 25 cm^2^ TC flask. Half of these aliquots were placed under 12:12 L:D at 21°C. The other half were enclosed in a film changing bag at 21°C to keep cells in constant darkness (D:D).

All aliquots were then left to rest overnight housed in the 21°C incubator in their respective conditions. Starting at ZT16 and continuing every 4 hours until ZT12, one culture from each condition (L:D or D:D) was retrieved at random under red light illumination and used to generate six 2 mL samples of cell suspension. Cells were centrifuged in the dark at room temperature for 5 minutes at maximum speed (12,800g) to pellet cells. Under red light, supernatant was removed and discarded with a micropipette; cell pellets were immediately flash frozen in liquid nitrogen.

After sample collection, total RNA was extracted from cell pellets using Trizol (ThermoFisher #15596026) following the manufacturer’s protocol, using glycogen as a carrier. RNA samples were treated with Turbo DNase (ThermoFisher #AM2238) to remove any contaminant DNA, then quantified using the Qubit HS RNA kit (ThermoFisher #Q32852) and assessed for integrity using the TapeStation 2200 (Agilent). Samples were submitted to The Bauer Core Facility at Harvard University for RNAseq library preparation using polyA selection with the KAPA HyperPrep kit (Roche) and sequenced with 150 bp paired end reads across 2 lanes of a NovaSeq flow cell (Illumina) at ∼60 million reads per sample. Reads are deposited in the NCBI SRA under BioProject ID PRJNA1242842.

### Circadian gene cycling analysis

RNAseq data were pseudo-aligned to the *E. muscae* transcriptome using Salmon [v1.10.1] with [-l A --validateMappings] to calculate isoform-level TPMs ^82^. Gene-level TPM was obtained by summing gene-specific isoform TPMs. Circadian cycling genes were identified using the meta2d function in the R package Metacycle ^47^, integrating the JTK_Cycle and LS methods, with default parameters. Genes with a Benjamini-Hochberg q-value below 0.05 were considered putative cyclers.

### Search for putative homologs to White Collar-1 and White Collar-2

To identify putative White Collar candidates, the *Neurospora crassa* White Collar-1 and-2 sequences were searched against the *E. muscae* genome using BLASTp using default settings. The resulting candidate proteins were assessed using PFAM ^83^ and HMMER ^84,85^ to identify protein domains and model protein homology to known white collar 1 and 2 sequences using default settings. The predicted *E. muscae* White Collar 1 candidates were required to contain PAS_9, PAS_3 and ZnF_GATA domains to be accepted as putative White Collar 1 proteins ^86,87^. The predicted White Collar 2 candidate was similarly assessed using PFAM and phmmer, and required to contain PAS_3 and ZnF_GATA domains to be accepted as a putative *E. muscae* White Collar 2 protein ^88^.

## Key resources table

| Reagent Type (species or resource) | Designation | Source or reference | Identifiers |
| --- | --- | --- | --- |
| genetic reagent (*D. melanogaster*) | Canton-S | <sup>31</sup> | Canton-S WF |
| genetic reagent (*D. melanogaster*) | per[01] | Bloomington Drosophila Stock Center | BDSC:80928 |
| genetic reagent (*D. melanogaster*) | per[30] | Bloomington Drosophila Stock Center | BDSC:63136 |
| genetic reagent (*D. melanogaster*) | cry[b] | Bloomington Drosophila Stock Center | BDSC: 80921 |
| genetic reagent (*D. melanogaster*) | per[s] | Bloomington Drosophila Stock Center | BDSC: 80919 |
| genetic reagent (*D. melanogaster*) | cry[02] | Bloomington Drosophila Stock Center | BDSC: 86267 |
| genetic reagent (*D. melanogaster*) | norpA | <sup>89</sup>; Gift from Dr. Kyobi Skutt-Kakaria | NorpA[p24] |
| genetic reagent (*D. melanogaster*) | CHO | <sup>43</sup>; Gift from Dr. Stan Lazopulo | CHO |
| strain (*Entomophthora muscae*) | *Entomophthora muscae* | <sup>33</sup> | ARSEF #13514 |

## Supplementary information

Supplemental File S1. MetaCycle results for *in vitro E. muscae* time course.

Supplemental File S2. Annotations for predicted cycling genes (FDR = 5%).

---

**Figure 1-S1.**
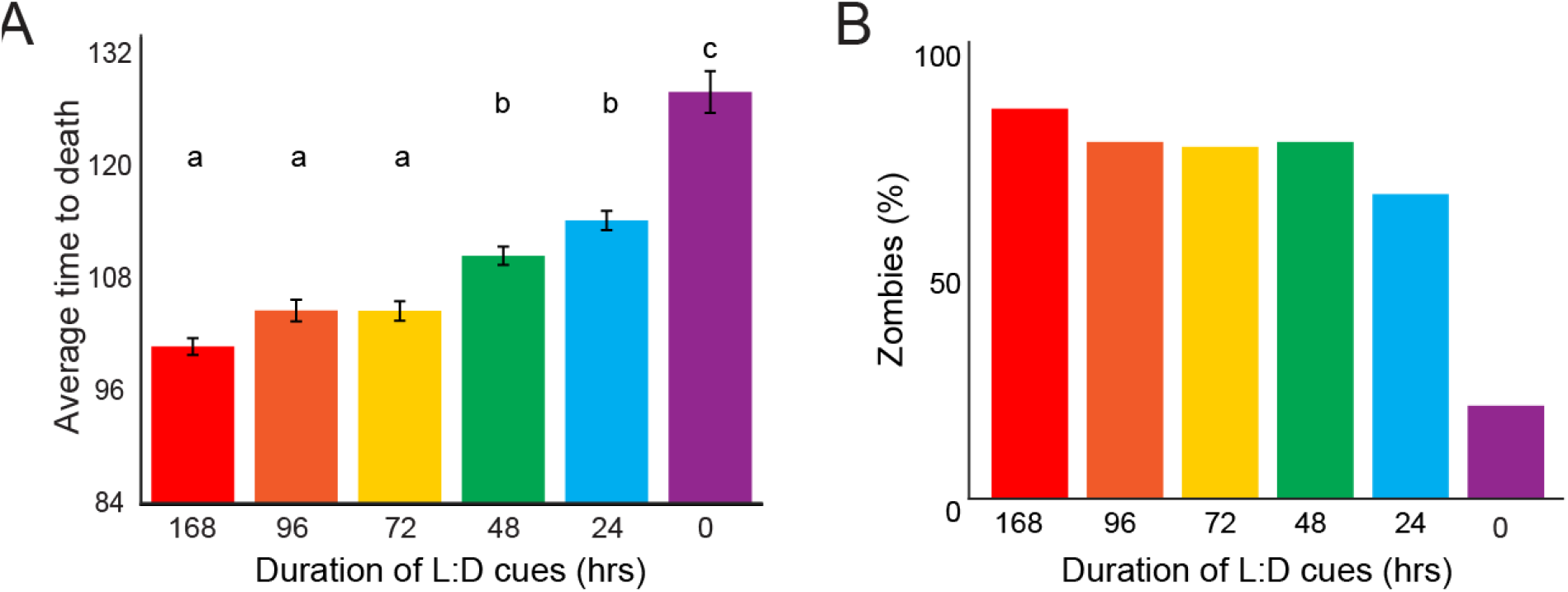
Observations in WT flies exposed to *E. muscae*. A) Mean time until death for WT (Canton-S) flies exposed to *E. muscae* over a range of lighting conditions. Error bars are SEM. Letters above bars indicate significance groups (p<0.05) as determined by comparisons (two-tailed t-test) for each experiment pair. B) Percentage of *E. muscae*-exposed flies that became zombies in Figure 1A.

**Figure 3-S1.**
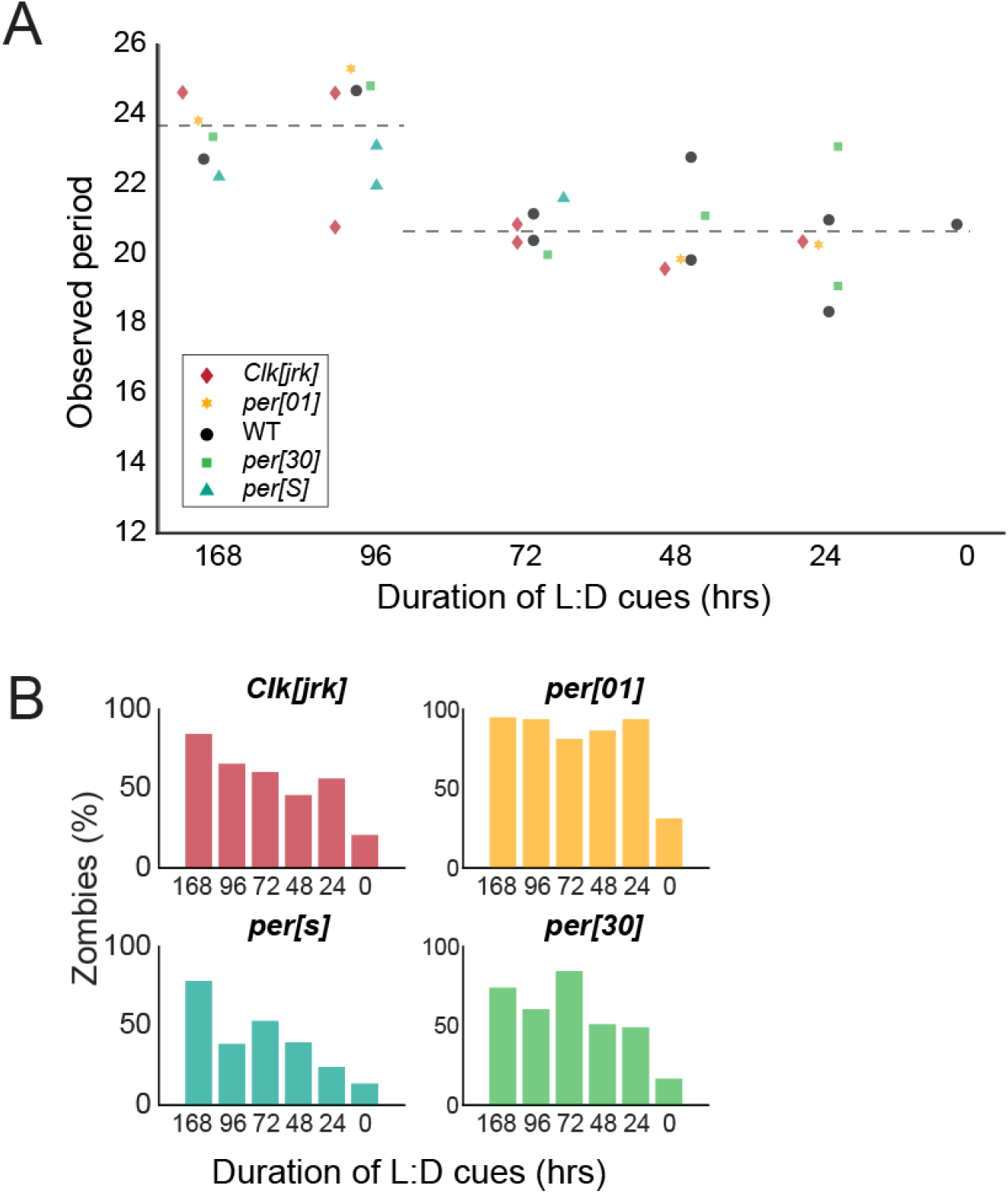
Observations in circadian mutants exposed to *E. muscae*. A) Observed death period lengths (in hours) for flies in Fig. 3B-G. For every day in which at least five flies were observed to die, the mean time of death was calculated; the difference between mean times of death across consecutive days is plotted. Dashed line at left reflects mean observed period for experiments where L:D cues were provided for experiments in which flies had at least one death day where L:D cues were provided (168 and 96 hours after *E. muscae* exposure; 23.7 hours); at right, mean for all experiments where flies were housed in D:D across all days of death (20.6 hours). Two-tailed t-test shows significant difference between L:D and D:D periods (p=5.2208e-05 excluding Cs WF periods, p=9.6136e-07 including Cs WF periods). B) Percentage of exposed flies that died and sporulated following *E. muscae* infection for flies in Fig. 3B-G.

**Figure 4-S1.**
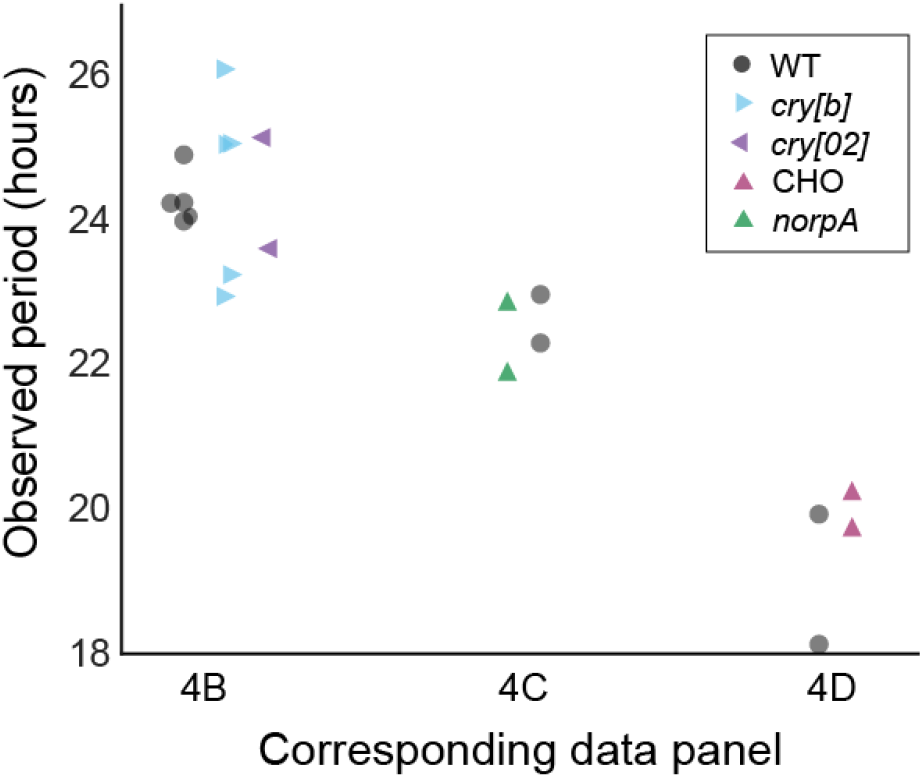
Observed death period lengths (in hours) for experiments in. Figure 4. For every day in which at least five flies were observed to die, the mean time of death was calculated and plotted above. Data from all replicates (not just representative experiments shown in Figure 4) were used. Mean observed period in experiments where the system could detect light (4B,4C) = 23.9; where the system could not detect light (4D) = 19.5.

**Figure 5-S1.**
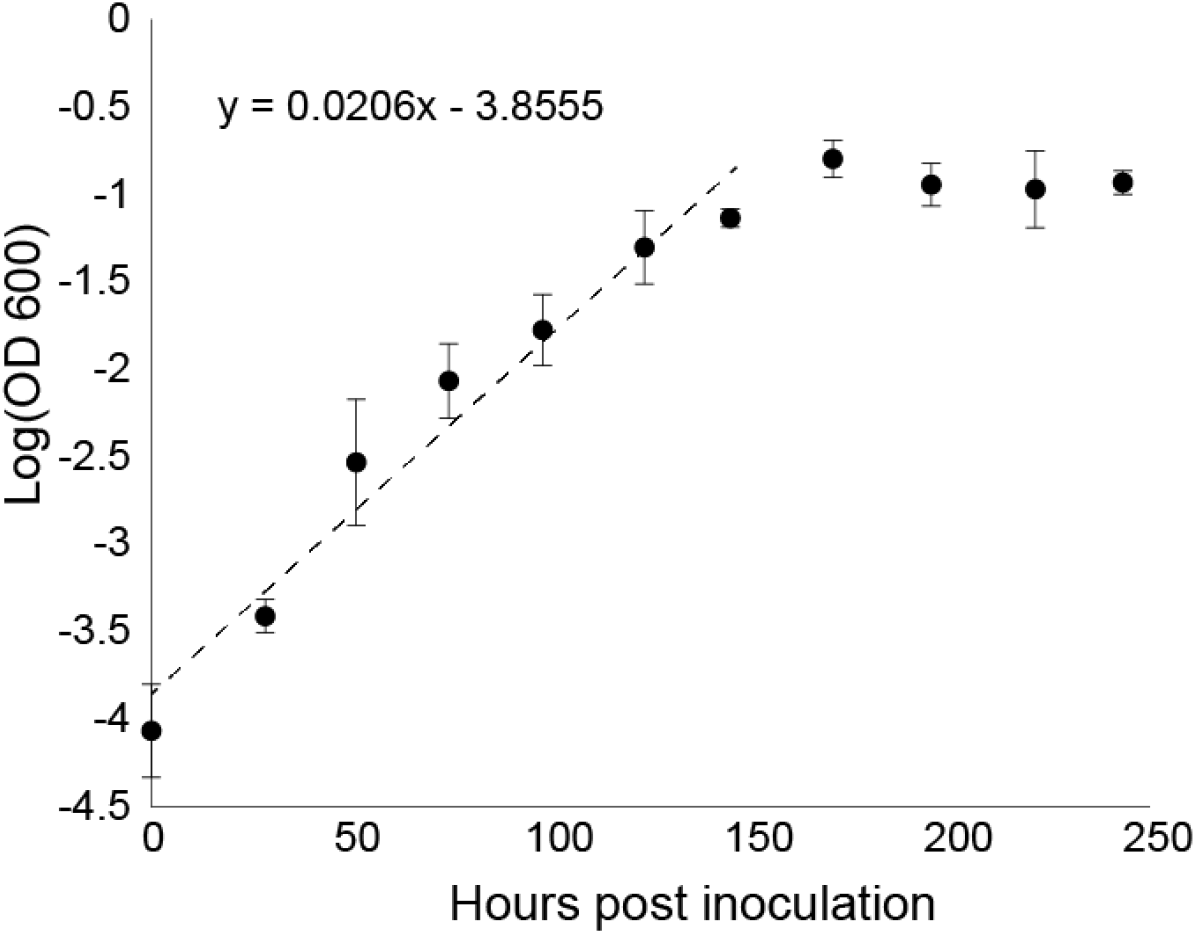
*In vitro E. muscae* growth curve at room temperature. *E. muscae* was at room temperature; OD600 was measured every ∼24 hours over 11 days (see Methods). Each dot represents the mean value of the triplicate cultures measured in triplicate; error bars show standard deviation between these values. Equation for line of best fit in linear region shown above; doubling over this time occurs every 30 hours on average.

**Figure 5-S2.**
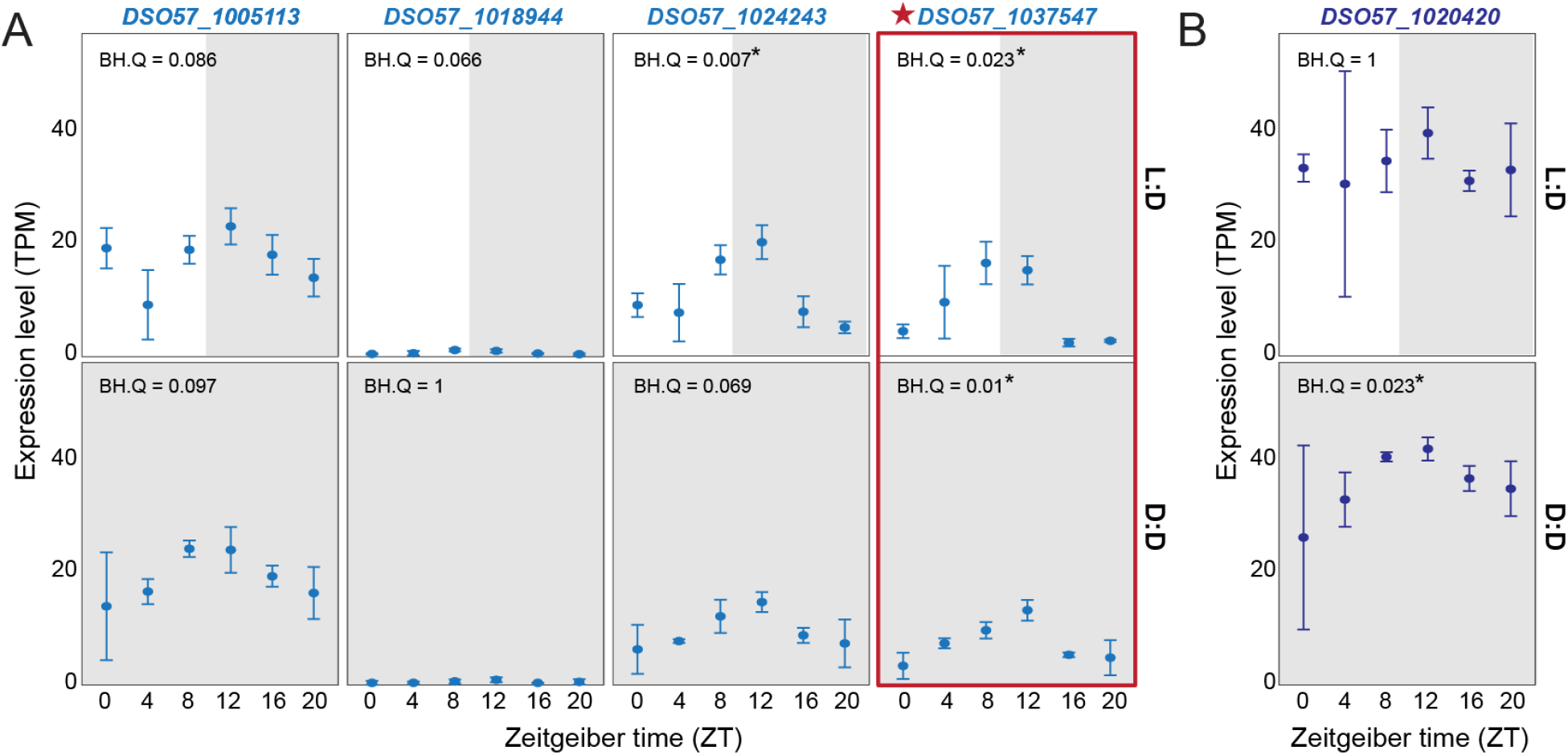
Expression patterns of *N. crassa* circadian homologs. Expression (in transcripts per million [TPM]) for A) each *white collar-1* homolog and B) the one *white collar-2* homolog across samples. Benjamini Hochberg Q (BH.Q) value is noted for each gene. BH.Q values below 0.05 are marked with an asterisk. To be considered a cycler (for Figure 5), BH.Q needed to be less than 0.05 across both L:D and D:D conditions. The one gene whose transcription pattern meets these criteria (*DSO57_1037547*) is outlined in red.

**Table S1.**
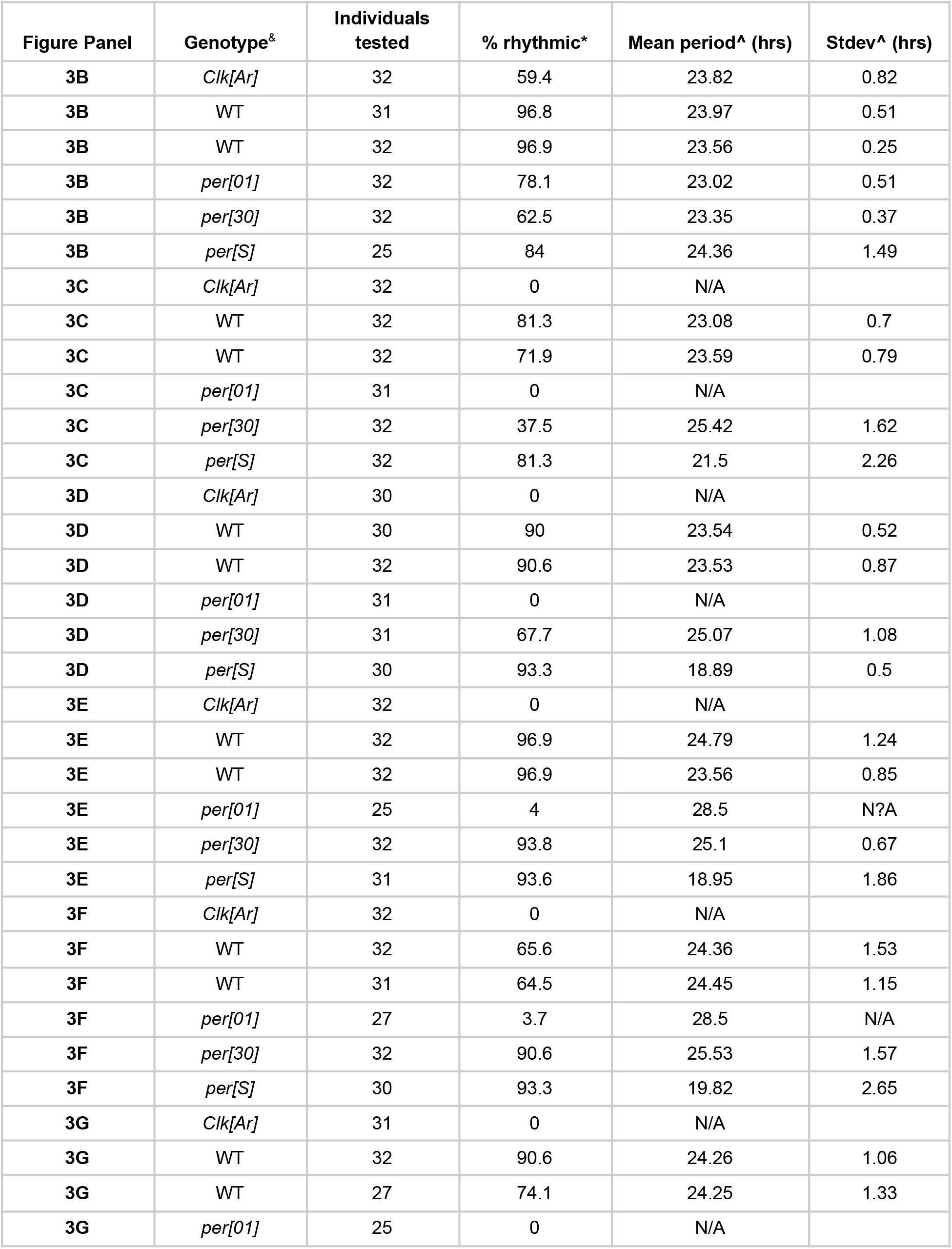
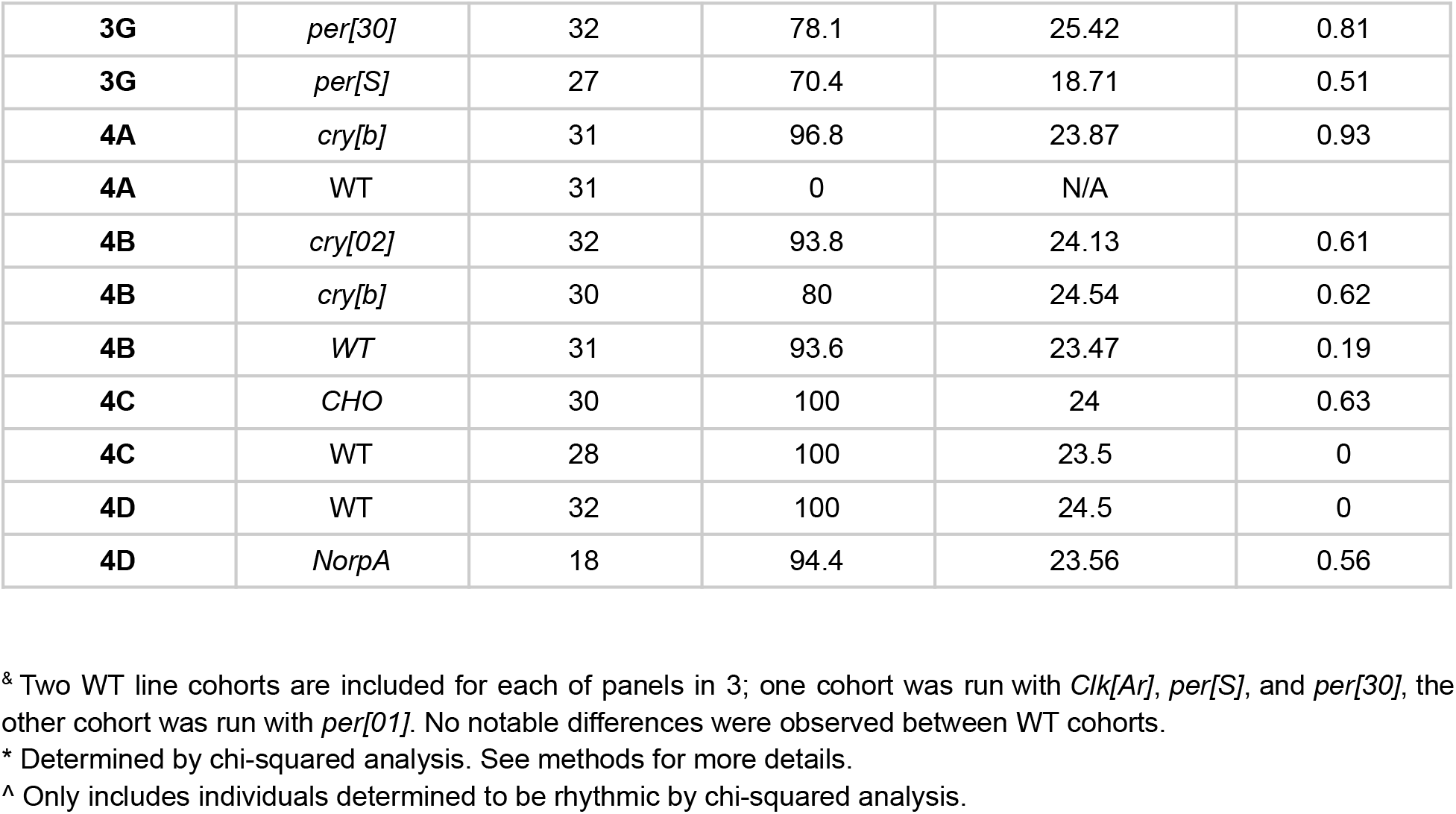
Periodogram analysis for uninfected controls run alongside exposed individuals.

**Table S2.**
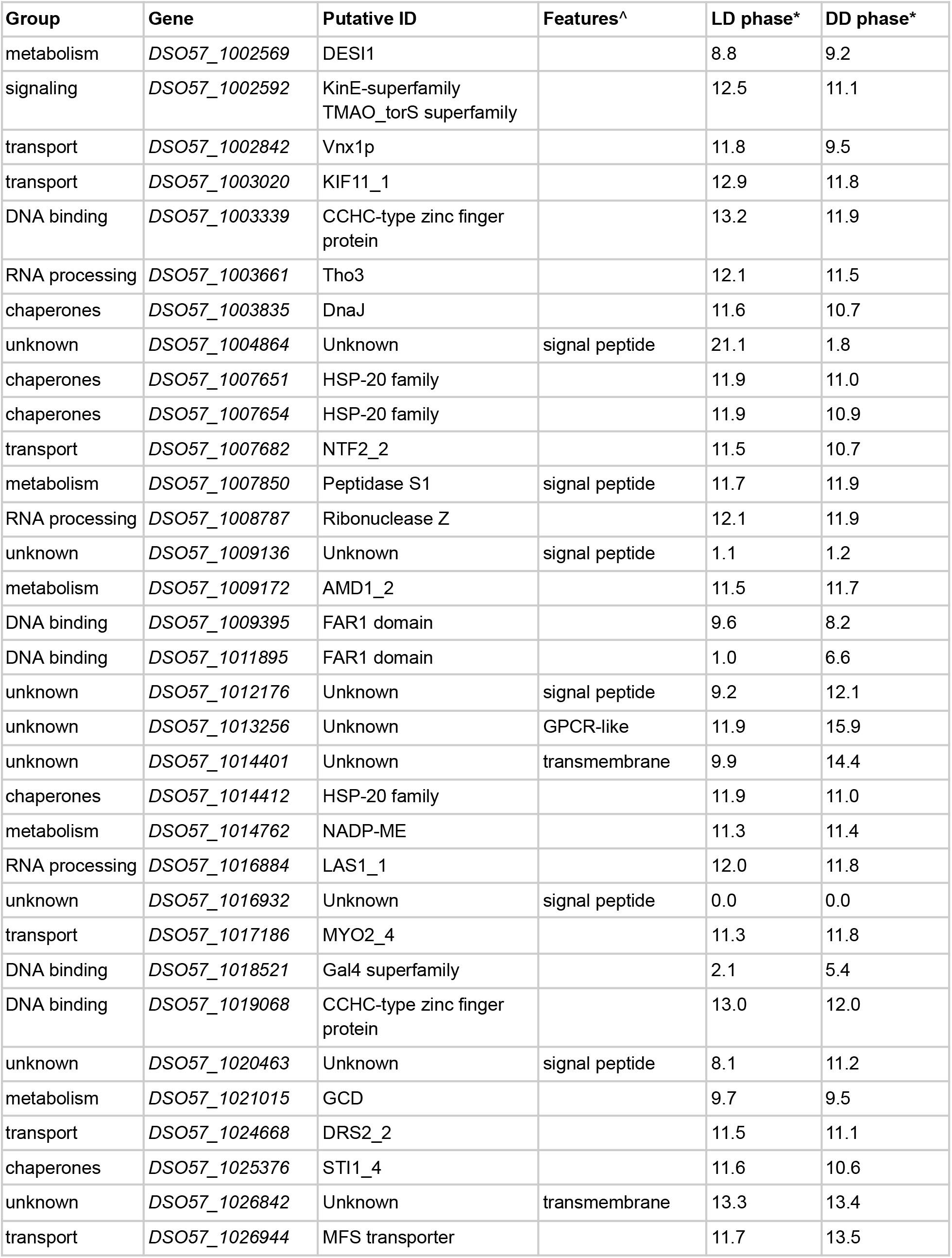
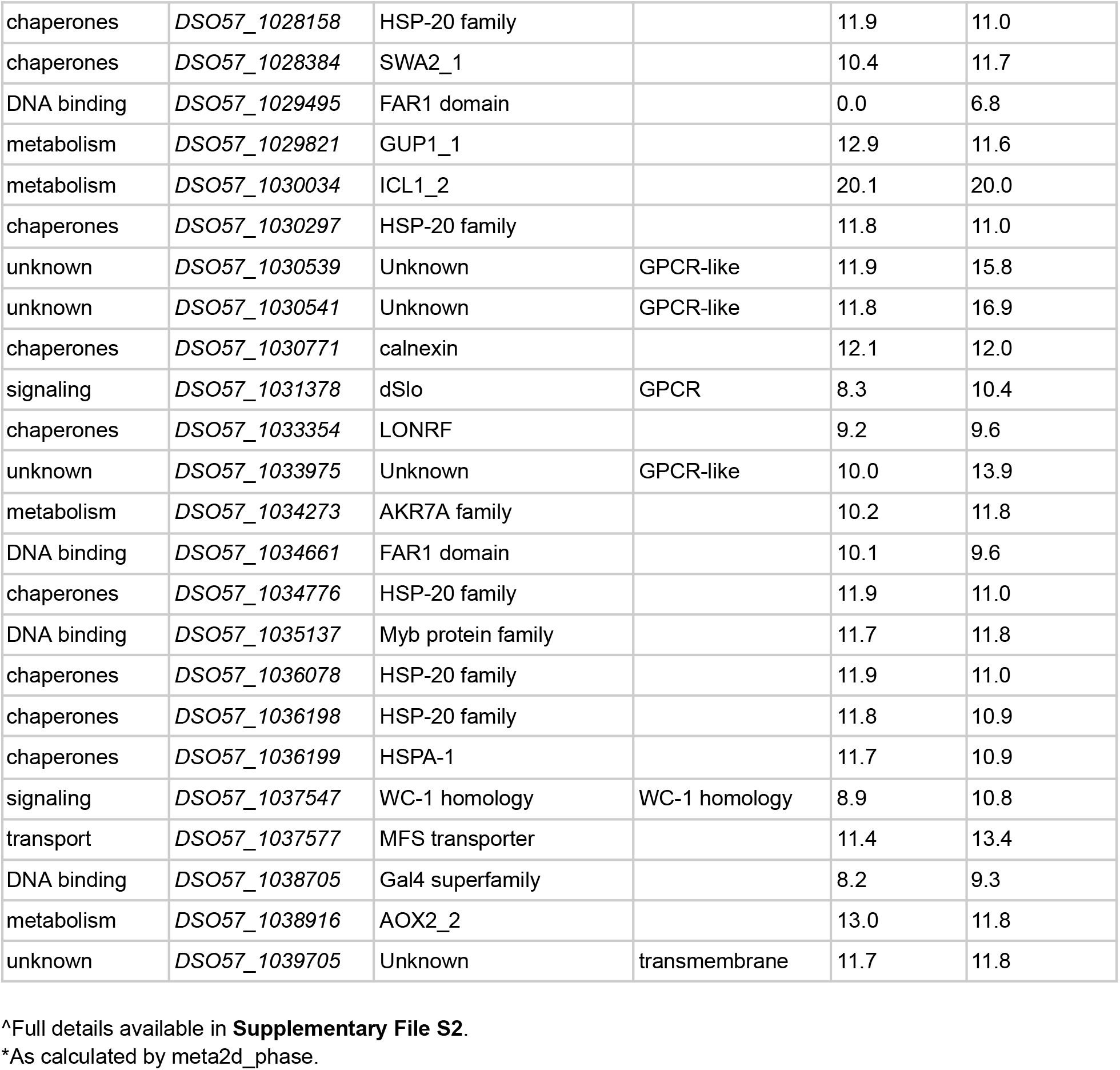
Functional groupings and phasing of cycling genes.

